# An explainable AI latent space of brain dynamics reveals a cerebello-prefrontal signature of schizophrenia symptoms

**DOI:** 10.64898/2026.08.25.746991

**Authors:** Moritz Bonhoeffer, Paolo Muratore, Mackenzie Weygandt Mathis, Indrit Bègue

## Abstract

Schizophrenia presents with several partially independent symptom dimensions, including positive symptoms, negative symptoms, and cognitive impairment; yet no neuroimaging framework has provided individual-level markers of symptom severity that remain anatomically interpretable. Here, we present an interpretable AI-based framework that addresses this gap by mapping high-dimensional resting-state rs-fMRI dynamics onto a low-dimensional latent manifold using self-supervised contrastive learning with a new attribution method to localize the highest decodable regions. Applied to two independent schizophrenia-spectrum cohorts, the label-free latent space supports individual-level prediction across clinical features of the disorder, including symptom severity and cognitive function. The attribution maps identify a disease-specific pathological footprint concentrated in prefrontal, posterior cerebellar and temporal areas that diverge from the manifold organization observed in healthy controls, which was dominated by auditory, limbic, and ventral-striatal circuits. These results establish an interpretable latent space framework for characterizing the distributed neural substrates of schizophrenia symptoms at the level of the individual patient, and provide an anatomically grounded route toward precision decoding of symptom severity.

---

Schizophrenia is a clinically heterogeneous disorder presenting with several partially independent symptom dimensions positive symptoms (hallucinations, delusions), negative symptoms (avolition, anhedonia, asociality, alogia, blunted affect) and cognitive impairment. Yet, despite decades of work, no neuroimaging framework reliably predicts symptom severity at the individual level across these dimensions. Clinical assessment currently relies on validated subjective rating scales; these instruments are coarse proxies for the neural processes they attempt to capture, summarizing behavior over weeks rather than indexing the mechanisms that generate it, and failing to capture individual variation in the underlying brain dynamics (1). There is a need for individual-level objective markers that characterize the distinct brain dynamics underlying symptom dimensions, which may in turn inform therapeutic symptom-specific strategies (2, 3). This need is most acute for negative symptoms that are the principal drivers of long-term functional disability (4), with no approved, mechanistically targeted intervention (5, 6). A framework capable of closing this gap across symptom dimensions is a prerequisite for the precision interventions that schizophrenia care currently lacks.

Resting-state functional MRI (rs-fMRI) has long been positioned as a candidate tool for this purpose, yet it has not fulfilled this promise in clinical practice (7). Traditional frameworks, such as mass univariate and seed-based connectivity, have provided foundational insights in schizophrenia. However, these methods are optimized primarily for group inference rather than individual prediction (6, 8). Thus, they may miss the distributed, multivariate patterns that characterize complex symptom dimensions, patterns potentially encoded in the high-dimensional, time-varying interactions of the whole-brain network (9). While approaches including ICA, PCA, and graph-theoretic methods capture more distributed structure, they frequently reduce brain dynamics to static summary metrics that discard the temporal geometry of neural activity (10). An alternative possibility is that clinically relevant information is encoded in the manifold geometry: specifically, in the shape of the low-dimensional manifold along which brain dynamics unfold over time (11, 12). To our knowledge, no existing framework simultaneously recovers this structure and renders it interpretable at the level of individual symptom severity.

This lack of interpretability is a critical barrier to mechanistic understanding, as it obscures the path from prediction to pathophysiology (13) to treatment. Identifiable self-supervised approaches offer a path forward by learning structure directly from the temporal organization of neural data. By leveraging the intrinsic consistency of brain dynamics rather than clinical ratings, these methods avoid the risk of overfitting to subjective scales while extracting high-dimensional representations that retain rich clinical information (14, 15). The CEBRA framework instantiates this principle, learning a low-dimensional latent manifold of neural time series through temporal contrastive objectives (16). Crucially, our extension to CEBRA introduces a Jacobian regularization term that renders the encoder analytically invertible without using auxiliary labels, enabling the construction of attribution maps that identify the contribution of individual brain regions to the learned representation (17). This combination is distinctive in the context of psychiatric symptom decoding: the latent space is learned without access to symptom labels, yet its constituent regions can be recovered analytically, and a single trained representation supports prediction of multiple clinical targets without retraining. Together, these properties allow the latent space to serve as a biologically grounded model of distributed brain function, providing a direct link between high-dimensional dynamics and the neurobiological substrates of the disease.

## Results

### A self-supervised encoder maps high-dimensional rs-fMRI dynamics onto a low-dimensional latent manifold

Our framework starts from preprocessed rs-fMRI signals and then, for each subject, constructs multi-dimensional time series by averaging voxel activity within atlas-derived regions of interest (ROI). We then leverage a new variant of CEBRA (16, 17) to build a low-dimensional latent representation of the rs-fMRI signal (Figure 1a, b). CEBRA learns a non-linear invertible encoder via temporal contrastive learning that maps the high-dimensional brain activity at each time point (for every subject) to a corresponding latent representation (see Methods). Importantly, we additionally incorporated two critical steps: Jacobian regularization for the encoder without using auxiliary labels, and an attribution method (17–19) that allows us to attribute the input (regional activity) to a particular latent dimension, which collectively is called xCEBRA-Time.

**Figure 1.**
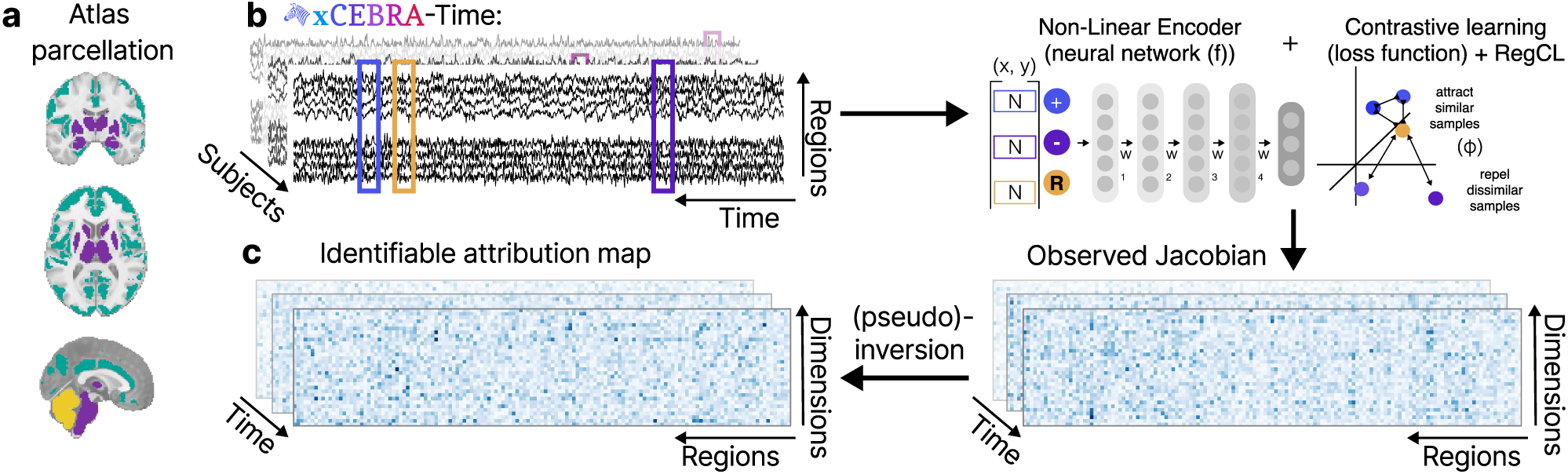
Interpretable framework for rs-fMRI based neural dynamics in schizophrenia. a: Patient and control rs-fMRI recordings are first preprocessed and ROI-averaged brain activity is assembled as a multi-dimensional time series. b: Our xCEBRA-Time framework is used to construct a low-dimensional representation of the signal (manifold) via a non-linear encoder. A positive sample is picked for each reference sample at a specific temporal distance, in our case 20 time points, within the same subject and session. Negative samples are drawn uniformly from the whole dataset. The non-linear encoder is then optimized to minimize the difference between the reference and the positive sample, while at the same time maximizing the difference between the reference and the negative samples. Additionally we introduce a Jacobian regularizer term, to be able to later invert the encoder. The value of the Jacobian regularizer is chosen to be at the highest value, without significantly increasing the final training loss. The CEBRA encoder (offset5-model) had 1024 hidden units, n=32 output dimensions, and was trained for 20k steps (see Methods). A separate decoder (a k-NN) is trained to leverage this 32-dimensional manifold to solve several regression tasks (e.g., symptom severity, see Figure 2).

Our proposed xCEBRA-Time provides the ability to map which parts of the rs-fMRI signal are relevant to shaping the manifold with theoretical guarantees under reasonable assumptions (see Methods). The resulting attribution maps reflect the construction that associates each input point (e.g., a single ROI at a given point in time) with a single scalar number quantifying its relative importance as inferred by the model. Intuitively, higher scores indicate a meaningful contribution to the overall construction, while small or otherwise negligible scores indicate how that particular input segment was not actively used.

Concretely, at every point in time *t*, each observation 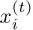 (rs-fMRI regional activation) is influenced by a collection of latent factors 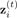 (hemodynamic response, neural activation), which, in turn, evolves into 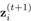 following some structure that defines how latent components interact (brain network dynamics). In these settings, the notion of attribution can be grounded in the ground truth process: we ask how the inferred attribution maps derived from recorded time-series data relate to the underlying data-generating process, i.e., the “ground truth map”, at each time step. This allows us to connect the attribution map to the causal structure of the data generating process. Namely, how (or if) each latent 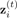 is related to our observations 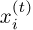. A core strength of the algorithm is a precise characterization of such relations: under reasonable assumptions, the inferred attribution maps are guaranteed to respect the (unobserved) underlying data-generating process (see Methods). This identifiability property is especially valuable in the medical setting, where attribution claims (e.g., if a given brain region is directly implicated in the process under study) are essential to inform subsequent interventions.

Lastly, after the unified encoder model is obtained, prediction is then performed by a separate k-Nearest Neighbors (kNN) regressor trained on the manifolds using nested cross-validation, with all data splits performed at the subject level so that frames from the same subject never appear simultaneously in training and evaluation sets.

### The latent space carries clinically meaningful structure across multiple symptom dimensions

We asked whether the xCEBRA-Time generated latent space is relevant to the clinical features of schizophrenia (Figure 2a). Across two independent cohorts, the unified manifold supported individual-level decoding-based prediction of every class of variable we tested (Figure 2b-f, Suppl. Figure S1). Specifically, for symptoms, we evaluated the Positive and Negative Syndrome Scale and found a stronger correlation for negative vs. positive symptoms (0.460 vs. 0.241, Figure 2b, c). While predictions for PANSS-positive scores were relatively weak (Combined: *r* = 0.241; Dataset 1: *r* = 0.214, Dataset 2: *r* = 0.166), PANSS-negative scores were decoded with higher accuracy across all subjects (Combined: *r* = 0.460; Dataset 1: *r* = 0.573; Dataset 2: *r* = 0.262), suggesting that the latent space captures a distinct neurobiological signature of a “deficit” syndrome, rather than a general marker of psychosis.

**Figure 2.**
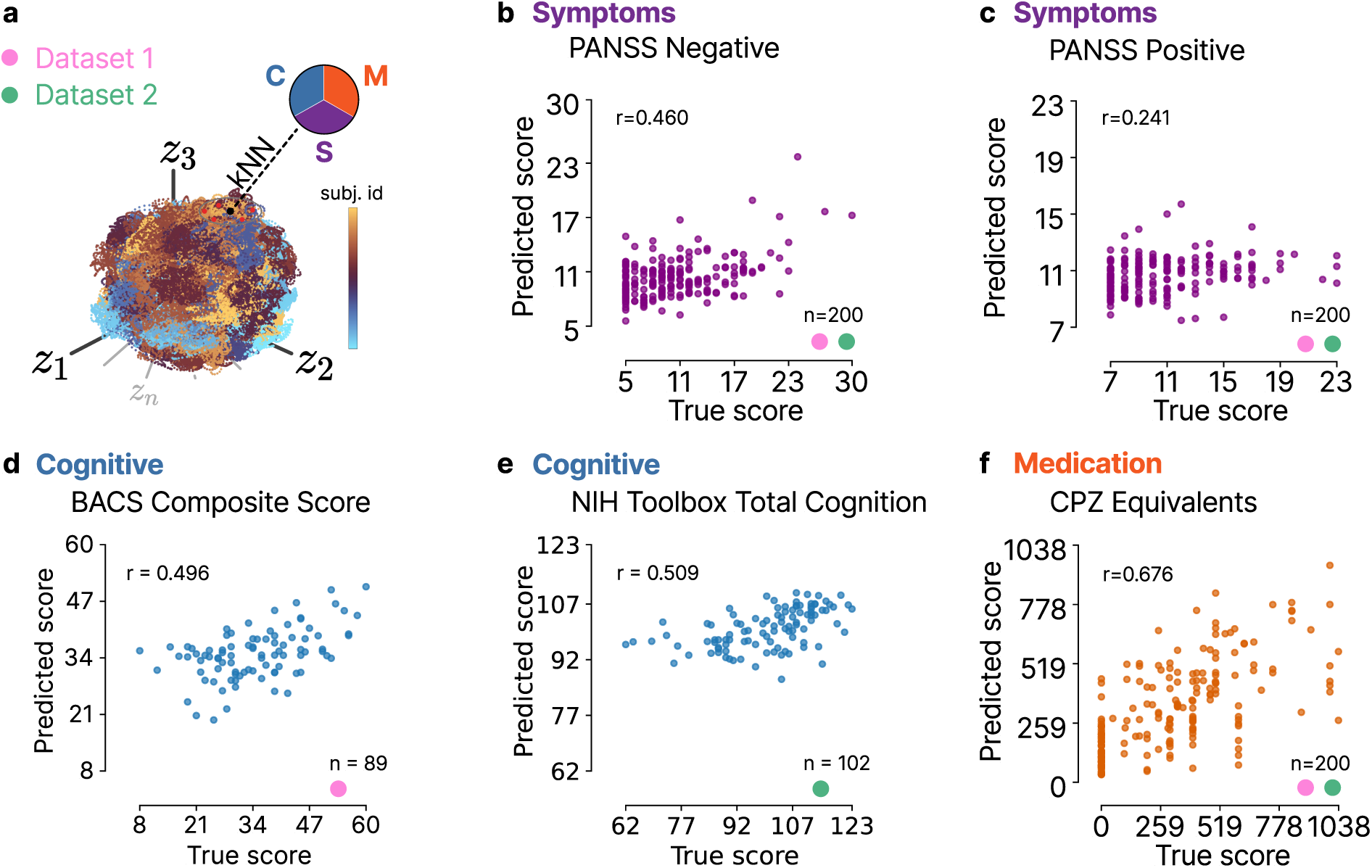
Latent representations support decoding of multiple clinical dimensions. a: We evaluate the decoding performance of symptoms (S), cognitive function (C), and medication (M) from the xCEBRA-Time manifold. For each k-NN decoder (i.e., in b–f) we report the number of datapoints (n), the Pearson correlation between predicted and true scores (r), its significance (p), and the explained variance (R2). b: PANSS-negative subscore (n = 200, r = 0.460, R2 = 0.209, p < 0.001). c: PANSS-positive subscore (n = 200, r = 0.241, R2 = 0.049, p < 0.001). d: Cognitive function measured by the BACS total score (n = 89, r = 0.496, R2 = 0.246, p < 0.001). e: Cognitive function measured by the NIH Toolbox Total Cognition composite score (n = 102, r = 0.509, R2 = 0.249, p < 0.001). f: Antipsychotic dose in chlorpromazine equivalents (n = 200, r = 0.676, R2 = 0.454, p < 0.001). Abbreviations: PANSS = Positive and Negative Syndrome Scale; BACS = Brief Assessment of Cognition in Schizophrenia, NIH Toolbox = NIH toolbox for assessment of neurological and behavioral function, CPZ = Chlorpromazine.

To test another dimension of impairment in schizophrenia, we tested whether cognitive function was decodeable from the xCEBRA-Time manifold. In Dataset 1 we were able to decode the BACS total score (20) as a marker of cognitive function with a correlation of *r* = 0.496. To further ensure this predictive capacity was not cohort-specific, we extended our decoding analysis to the Human Connectome Project for Early Psychosis (HCP-EP) dataset (Dataset 2), using the NIH total cognition composite score (21, 22), with the framework yielding comparable and robust accuracy (*r* = 0.509, Figure 2e). This confirms that the learned latent representations capture neural signatures of cognitive impairment across independent populations.

We further assessed whether the xCEBRA-Time manifold captured neural signatures of antipsychotic exposure. The decoder accurately predicted chlorpromazine (CPZ)-equivalent doses across the combined dataset (*r* = 0.676, Figure 2f), with robust performance maintained in independent cohorts (Dataset 1: *r* = 0.518; Dataset 2: *r* = 0.526). These collective results indicate that the xCEBRA-Time manifold effectively encodes the systematic influence of pharmacological treatment on distributed functional networks alongside disease-specific pathology.

Lastly, given the framework’s ability to decode both medication dose and symptom severity, we sought to determine whether the relationship between the latent space and symptom severity or cognitive function was confounded by antipsychotic exposure. We performed a partial correlation analysis between true and predicted scores for each of the clinical parameters (positive, negative symptoms as well as cognitive function), controlling for antipsychotic dosage in CPZ-equivalents. The association between latent representations and the predicted clinical and cognitive scores remained essentially unchanged (see Table S1), suggesting that xCEBRA-Time captures intrinsic pathological brain dynamics independent of medication load. To exclude head motion confounds, we repeated the partial correlation analyses controlling for mean framewise displacement (Table S1) and observed the same pattern.

### The latent space is driven by a disease-specific signature in the posterior cerebellar, prefrontal, and temporal cortices

We leveraged our method and model to subsequently identify the anatomical features driving the xCEBRA-Time manifold (which was trained with all control and patient data). We quantified the relative contribution of each region to the construction of the latent space, isolating the neural features most critical for symptom decoding. The highest cortical attribution scores localized to the bilateral middle temporal gyri and the frontal poles of the prefrontal cortex (Table S2). Subcortically, the brainstem emerged as a primary contributor, followed by the thalamus (Table S2). Notably, within the cerebellum, predictive weight was concentrated specifically in the posterior lobules (VI and VIIb) (Table S2). This anatomical profile suggests that the latent space is heavily influenced by regions associated with high-level cognitive integration.

To confirm that these regional attributions reflected pathological signatures rather than general features of neural dynamics during resting state, we performed a specificity analysis by training separate models on patient and healthy control cohorts (Figure 3a, b). The patient-only attribution map closely recapitulated the spatial topography of the primary combined model, centered on the aforementioned cerebello-temporo-prefrontal nodes (Figure 3a, Table S3). In contrast, the healthy control model exhibited a markedly different profile, driven primarily by sensory-processing regions including the planum polare, planum temporale, and Heschl’s gyrus (Figure 3b, Table S4). Moreover, the stability of these maps under systematic parameter sweeps was confirmed (Suppl. Figure S2). Then we computed the difference in the attribution maps between patients and controls (see Methods). Strikingly, we see that the largest changes are in the cerebello-temporo-prefrontal regions (Figure 3c, Table S5). Taken together, this divergence between patient and control maps underscores that the model’s sensitivity to negative symptoms is driven by a unique pathological footprint, characterized prominently by posterior cerebellar, temporal and prefrontal contributions that remain stable across independent cohorts (Figure 3a-c, Tables S2-5).

**Figure 3.**
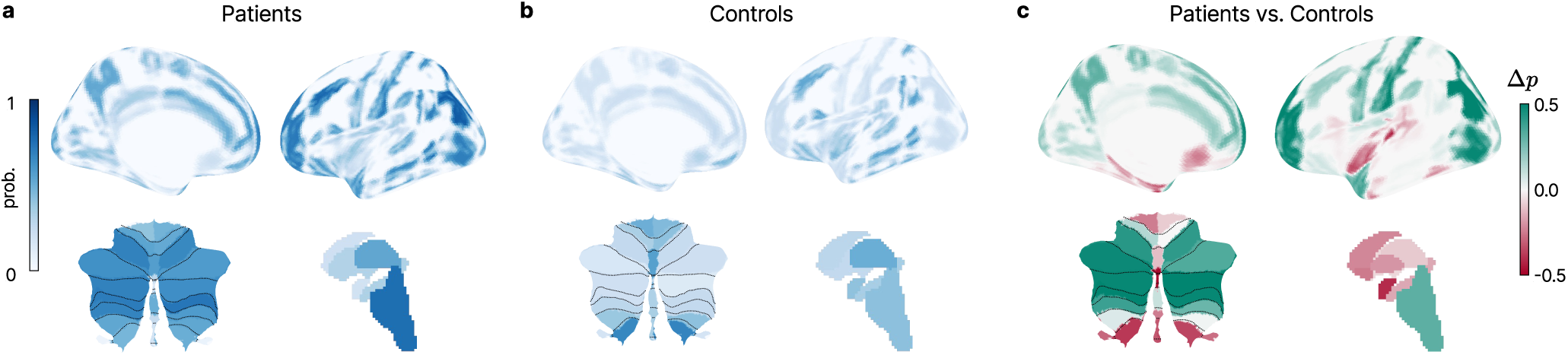
xCEBRA-Time attribution maps identify disease-specific brain regions. a: For three trained xCEBRA-Time models, we computed the inverse Jacobian matrix using Singular Value Decomposition. Prior to inversion, the absolute values of the Jacobian matrix were taken to capture the magnitude of influence. To establish a global measure of regional importance, the time-resolved matrices were averaged across all time points in the dataset, producing a single summary matrix (input dimensions × output dimensions) for each model. To correct for initialization inconsistency, the attribution maps of the three CEBRA models were Procrustes aligned. Because xCEBRA-Time can only discern between zero and non-zero elements, the aligned maps were binarized at z = 0 and subsequently averaged to form a single consensus attribution map. For visualization on a standard MNI atlas, this consensus map was averaged across the output dimensions to generate a single importance score (ranging from 0 to 1) for each region of interest (ROI). (a:) Reconstructed attribution map from an xCEBRA-Time model trained solely on the patient cohort. (b:) Reconstructed attribution map from a model trained exclusively on the healthy control cohort. (c:) The difference between the patient and control attribution maps after the respective attribution maps were mean-centered. Regions highlighted in green contribute disproportionately in patients, indicating disease-specific areas.

## Discussion

We set out to address a long-standing gap in biological psychiatry: the absence of objective, individual-level measures of symptom severity in schizophrenia. Using a new self-supervised contrastive framework (xCEBRA-Time), we learned a lowdimensional latent manifold of rs-fMRI dynamics that decoded symptom severity, cognitive function, and medication dose across two independent chronic schizophrenia and early-psychosis cohorts. Additionally—because the encoder is rendered analytically invertible—we localized the contributing signal to a stable anatomical footprint mapping onto higher-order associative networks (frontoparietal control and default mode), the cerebellar cognitive-affective system, and sensorimotor cortex. Three features distinguish this result from prior efforts: unlike case–control classifiers and connectivity-based predictive approaches, the latent space is learned without access to clinical labels; the decoding targets continuous symptom severity at the individual level; and the attribution map carries formal identifiability guarantees rather than the heuristic interpretability of post-hoc saliency methods.

Turning to the anatomical structure of this footprint, we find implicated regions from higher-order associative (bilateral middle temporal gyri, frontal poles, superior frontal gyri), cerebellar cognitive-affective (posterior cerebellar lobules VI, VIIb, Crus I/II, VIIIa), and sensorimotor systems (precentral and postcentral gyri). The divergence from the control manifold — whose organization was dominated by regions of the primary auditory network (Heschl’s gyrus, planum temporale, planum polare), the medial-temporal and limbic affective system (parahippocampal cortex, amygdala, hippocampus), the default-mode-linked cerebellar lobule IX and vermis, and the classical dopaminergic ventral-striatal targets (nucleus accumbens, pallidum, caudate) — further supports the interpretation that the patient footprint reflects a disease-specific reorganization of the latent geometry rather than a generic property of brain dynamics. Notably, several of these control-anchored networks (auditory, limbic, ventral-striatal) are the very circuits classical accounts of psychosis emphasize as substrates of positive symptoms, suggesting that these circuits continue to structure intrinsic dynamics in the healthy brain but are no longer load-bearing in the patient manifold. Consistent with this dissociation, that the footprint is recovered by a manifold sensitive to negative but not positive symptoms supports the view that symptom dimensions in schizophrenia arise from partially distinct neural networks. One interpretation is that negative symptoms reflect a comparatively stable reorganization of intrinsic dynamics that resting-state contrastive learning is well suited to capture, whereas positive symptoms are more state-like and event-driven and may be better indexed by task or symptom-capture paradigms — a state/trait dissociation our framework’s asymmetric decoding directly implies but does not test.

The anatomical footprint we identify aligns with independent bodies of prior work. First, it converges with a broader connectivity-based literature that has independently implicated prefrontal and posterior cerebellar regions as prominent nodes of schizophrenia-related network alterations, alongside disrupted contributions from auditory, limbic, and ventral-striatal circuits (23–29).Second, it aligns with an emerging literature linking the cerebellum and its midbrain dopaminergic connections to reward, motivation, and social behavior — and specifically to cerebellar–ventral tegmental area dynamics in apathy (30–33) – a link that our findings further support by identifying the brainstem itself as a load-bearing contributor to the patient manifold. That an unsupervised manifold recovers these patterns from a fundamentally different analytic principle — the temporal geometry of intrinsic dynamics rather than pairwise coupling, and without access to clinical labels — argues that the network reorganization is a genuine property of the data rather than a method-specific artifact. This is further supported by the stability of the maps: patient-only and combined-cohort attributions converged on the same anatomical footprint, and the maps remained stable under a systematic sweep of architectural hyperparameters. Beyond confirming known structure, our analysis sharpens the picture in two ways: the prefrontal contribution localized specifically to the bilateral frontal poles rather than to prefrontal cortex more broadly — a regional specificity that connectivity-based analyses have not previously resolved, and one consistent with the frontal pole’s role in prospective cognition and goal maintenance (34) — and the dissociation between patient and control manifolds proved categorical rather than graded, with over twenty regions showing between-cohort differences greater than |Δ| *>* 0.30.

Importantly, our framework generates several falsifiable predictions for future work. First, longitudinal studies could test whether the latent signature tracks symptom change within the same patient and predicts treatment response, extending the framework from concurrent decoding toward prognostic utility — with the further prediction that the negative-symptom signal, which was weaker in the early-psychosis cohort, should strengthen longitudinally as the syndrome becomes chronic. Second, baseline manifold coordinates should carry predictive information about response to interventions targeting the negative-symptom dimension. Third, if the posterior cerebellar and prefrontal nodes are load-bearing rather than incidentally expressed, non-invasive stimulation targeted to these regions should measurably shift the manifold and, if the manifold is symptom-relevant, correspondingly alter negative-symptom severity. This prediction is of particular translational interest because the posterior cerebellum is accessible to non-invasive transcranial magnetic stimulation, and recent electric-field modeling work has begun to map stimulation targets for schizophrenia (3); an interpretable biomarker that nominates a specific, anatomically grounded node could help shift the search for negative-symptom therapeutics toward hypothesis-driven targeting. Testing the sharper prediction — dissociable effects of stimulating the posterior lobules (VI, VIIb, Crus I/II, VIIIa) versus lobule IX — will require higher-resolution stimulation protocols such as low-intensity focused ultrasound (LIFU). Fourth, if the signature reflects a general motivational-deficit dimension rather than a schizophrenia-specific marker, it should generalize transdiagnostically to apathy in major depression, Parkinson’s disease, or frontotemporal dementia. More broadly, because the method is agnostic to data modality and depends only on the temporal organization of the signal, the same identifiable-manifold approach could be applied to other disorders and recording modalities where individual-level, mechanistically interpretable markers are needed.

In summary, we introduce a self-supervised, interpretable framework that decodes negative-symptom severity in schizophrenia at the individual level across two independent cohorts, without access to clinical labels. Our approach localizes the underlying signal to a stable, disease-specific footprint spanning higher-order associative, cerebellar cognitive-affective, and sensorimotor systems, with the bilateral frontal poles and posterior cerebellar lobules emerging as the most distinctive contributors. Because the attribution carries formal identifiability guarantees rather than heuristic interpretability, this footprint is not only predictive but inspectable — a link from prediction to anatomy that current deep-learning biomarkers do not provide. We suggest this offers a template for the kind of transparent, anatomically grounded biomarker that has so far eluded computational psychiatry.

## Acknowledgments

We acknowledge funding from the Leenaards Foundation and PRD 22-2020-I (awarded to IB), and the SNSF Grant No. 10.000.950 (awarded to MWM).

## Author Contributions Statement

Conceptualization: IB, MWM, MB, PM Methodology: MB, PM, MWM, IB Investigation: MB, PM Writing-Original Draft: MWM, IB Writing-Editing: MB, PM, MWM, IB Funding Acquisition: MWM, IB

## Declaration of interests

M.W.M. is the co-inventor of CEBRA, and its use for brain decoding is covered under US Patent No. US12499131B2. The other authors declare no conflicts of interest.

## Online Methods

### Dataset description

The training data were aggregated from two independent datasets, combining longitudinal and cross-sectional functional MRI (rs-fMRI) recordings.

### Overview

#### Dataset 1: In-house Cohort

The data were obtained from clinical cohorts approved by the Geneva Ethics Committee (BASEC IDs: 2017-01765, 2020-02169). This initiative was designed to find markers for motivational negative symptoms. Data collection was conducted at the community outpatient center of the University Hospital of Geneva, Switzerland.

The clinical cohort consists of clinically stable outpatients with a confirmed schizophrenia spectrum diagnosis, assessed via the Mini International Neuropsychiatric Interview. Clinical stability was defined by no hospitalizations or medication changes in the preceding three months. A cohort of matched healthy controls with no personal history of Axis-I disorders, psychotropic drug use, or a family history of psychotic disorders was acquired. Important exclusion criteria included secondary sources of negative symptoms (e.g., florid psychosis, extrapyramidal effects, or comorbid major depressive episodes), active substance use, or the use of benzodiazepines exceeding 1 mg of lorazepam equivalents. A full description of the dataset can be found here (32).

In addition to the neuroimaging data, extensive clinical, behavioral, and cognitive testing of the subjects was performed longitudinally, and antipsychotic medication was measured in risperidone equivalents. From an initial screening of 427 individuals, the baseline cohort consisted of 146 subjects, including 90 patients with schizophrenia spectrum disorder and 56 healthy controls (with a subset of 65 subjects returning for a 3-month follow-up). After exclusion of subjects with excessive movement, no cerebellum in field of view, interrupting the study or epileptic seizure we were left with 66 patients (25 with two sessions) and 49 healthy controls (19 with two sessions).

#### Dataset 2: HCP-EP Cohort

The data for the second dataset were obtained from the Human Connectome Project for Early Psychosis (HCP-EP) database, available through the National Institute of Mental Health (NIMH) Data Archive (NDA). The HCP-EP is a multi-site initiative designed to map macroscopic human brain circuits and their relationship to behavior in individuals during the early stages of psychotic illness. Data collection was conducted across four sites: Brigham and Women’s Hospital (coordinating center), Beth Israel Deaconess Medical Center, McLean Hospital, and Indiana University.

The dataset includes individuals aged 16 to 35 years. The clinical cohort consists of outpatients within the first five years of onset of psychotic symptoms. Patients with bipolar disorder or major depression with psychotic features, schizophrenia, schizoaffective disorder, or schizophreniform disorder, diagnosed according to DSM-5 criteria, were included. A cohort of demographically matched healthy controls with no personal history of psychotic disorders or first-degree relatives with a schizophrenia spectrum disorder is also included. Important exclusion criteria were substance-induced psychosis, IQ < 70, or subjects with an active medical condition that affects brain or cognitive functioning. In addition to neuroimaging data, extensive clinical and cognitive testing of the subjects was performed, medication was measured in CPZ-equivalents. An extensive description of the data set can be found in the reference manual (35). After exclusion of patients without neuroimaging or clinical data and those with excessive head-movement during the rs-fMRI scan, this cohort consisted of 143 subjects, including 110 patients with early psychosis and 33 healthy controls.

### Acquisition details

#### Dataset 1: In-house Cohort

All imaging data were acquired using a Siemens 3T Magnetom Prisma scanner equipped with a standard 64-channel head coil. The structural acquisition parameters included a repetition time (TR) of 2200 ms, an echo time (TE) of 2.96 ms, and a flip angle of 9°, utilizing anterior-to-posterior phase encoding without fat suppression. Data were collected with a field of view (FOV) of 256 mm and a slice thickness of 1.0 mm, yielding an isotropic spatial resolution of 1.0 mm³.

Resting-state functional MRI (rs-fMRI) scans were acquired over a single run lasting 9.8 minutes. Functional images were collected utilizing an acceleration mode with a parallel imaging factor of 6. The specific acquisition parameters included a repetition time (TR) of 1000 ms, an echo time (TE) of 32.0 ms, a flip angle of 50°, and anterior-to-posterior phase encoding. Data were acquired with a field of view (FOV) of 224 mm, featuring a 2.0 mm isotropic spatial resolution. Further details can be found here (3)

#### Dataset 2: HCP-EP Cohort

All data were acquired using Siemens 3T Magnetom Prisma scanners. The McLean Hospital site employed a 64-channel head and neck coil with the neck receivers deactivated, while the other sites used standard 32-channel head coils.

High-resolution anatomical T1-weighted images were acquired using a 3D magnetization-prepared rapid gradient-echo (MPRAGE) sequence based on the Human Connectome Project Lifespan protocol. The structural acquisition parameters in-cluded a repetition time (TR) of 2400 ms, an echo time (TE) of 2.22 ms, an inversion time (TI) of 1000 ms, and a flip angle of 8°. Data were collected with a field of view (FOV) of 256 mm, yielding a sub-millimeter isotropic spatial resolution of 0.8 mm. Following structural imaging, participants completed a total of four rs-fMRI runs within a single one-hour imaging appointment, with each run lasting 5 minutes and 28 seconds. Functional images were collected utilizing blood oxygenation level-dependent (BOLD) contrast via a gradient-echo echo-planar imaging (EPI) sequence, which incorporated a multiband acceleration factor of 8. The specific acquisition parameters were as follows: a field of view measuring 208 mm × 180 mm × 144 mm and 72 total slices. The data featured a 2 mm isotropic spatial resolution and a repetition time (TR) of 800 ms.

### rs-fMRI preprocessing

Dataset 1 was preprocessed utilizing SPM12. The structural and functional data underwent standard realignment and coregistration. Next, the structural images were segmented to generate tissue probability maps, which were then used to non-linearly normalize the functional data into standard MNI space. To ensure data integrity, visual quality control inspections were conducted after each processing stage.

Dataset 2 was processed using the standard Human Connectome Project (HCP) minimal preprocessing pipeline. This workflow included correcting for gradient non-linearities, *B*_0_ magnetic field distortions, and participant motion (using a single-band reference image), alongside structural registration. To minimize interpolation errors, all spatial transformations were concatenated and applied in a single step to resample the data into 2 mm MNI space. The rs-fMRIs were masked for non-brain tissue and intensity-normalized by a single scaling-factor, as well as minimally high-pass filtered (0.0005 Hz). After this and Independent Component Analysis was run utilizing MELODIC. Confounding components as classified by FIX, were removed from the data.

Identical voxelwise timeseries processing was applied to both datasets. First, the initial 10 seconds of each scan were discarded to allow for scanner magnetization equilibration. Next, to avoid partial volume effects, restrictive masks from the DPARSFA toolbox were used to extract the average white matter (WM) and cerebrospinal fluid (CSF) signals. The global signal was also extracted using voxels with a probabilistic grey matter density greater than 0.025.

Finally, the individual voxelwise timeseries were detrended and cleaned using a nuisance regression model that included the following confounders:

1. Linear and quadratic trends.
2. The average WM, CSF, and global signals.
3. A discrete cosine transform (DCT) basis acting as a high-pass temporal filter (frequency cutoff: 0.01 Hz).
4. The six standard head movement parameters derived from spatial realignment.
5. Spike regressors to scrub individual volumes corrupted by excessive motion, defined by a framewise displacement (FD) threshold exceeding 0.5 mm.

### rs-fMRI timecourse extraction

To reduce the dimensionality of the voxel-wise rs-fMRI data, we performed a Region-of-Interest (ROI) based time series extraction. Regional time courses were extracted using a composite parcellation scheme derived from three standard neuroanatomical atlases:

1. Harvard-Oxford Cortical Structural Atlas
2. Harvard-Oxford Subcortical Structural Atlas: To only include meaningful BOLD signal and avoid redundancy, we excluded ROIs corresponding to the ventricles, white matter, and global cortical signal (to prevent overlap with the cortical atlas).
3. Diedrichsen Cerebellar Atlas The time series for each defined ROI were computed by averaging the BOLD signal across all voxels within the region mask.

### In-Scanner Head Motion Quantification

To ensure that the predictive performance of our models was driven by true neurobiological variance rather than being a spurious byproduct of in-scanner head motion, we quantified participant movement using Mean Framewise Displacement (MFD). Motion estimation was based on the six realignment parameters (three translational and three rotational). These parameters were generated using standard SPM preprocessing for Dataset 1, and using FSL’s MCFLIRT (as part of the HCP minimal preprocessing pipeline) for Dataset 2. Framewise displacement (FD) was calculated for the time points as the sum of the absolute frame-to-frame changes across all six parameters. Rotational changes were converted to millimeters assuming a standard 50 mm head radius. The FD of the first retained volume was set to zero. The MFD was then calculated as the average displacement across all retained volumes for each subject. This MFD metric was subsequently utilized in partial correlation analyses to regress out the effects of head motion, confirming that our models’ predictive accuracies were independent of movement-related artifacts.

### Clinical and cognitive scores

#### Positive and Negative Syndrome Scale (PANSS)

Symptom severity was evaluated using the Positive and Negative Syndrome Scale (PANSS) (36). The PANSS is a semistructured clinical interview where each item is rated on a 7-point Likert scale ranging from 1 (absent) to 7 (extreme). For the purposes of this study, we utilized only the Positive and Negative subscales. In accordance with guidelines from the European Psychiatric Association (EPA), negative-symptom severity was quantified using the total sum score only of the items N1,N2,N3,N4,N6 (Marder factor of negative symptoms) (5). Positive symptom severity was quantified using the total sum score of all positive items (P1-P7).

#### Brief Negative Symptom Scale (BNSS)

To provide a granular and targeted assessment of motivational deficits, we administered the Brief Negative Symptom Scale (BNSS) (37) in our in house dataset (Dataset 1). Items on the BNSS are anchored on a 7-point scale ranging from 0 (normal symptom manifestation) to 6 (extremely severe deficit). In this study, we specifically isolated and analyzed the Apathy subscale to directly evaluate reductions in self-initiated, goal-directed behavior and internal motivation (sum of Avolition, Anhedonia and Asociality items).

#### Brief Assessment of Cognition in Schizophrenia (BACS)

To evaluate cognitive functioning, we utilized the BACS (20) in our in house dataset. The BACS assesses six cognitive domains associated with schizophrenia — verbal memory, working memory, motor speed, verbal fluency, attention and processing speed, and executive function. We used the BACS composite score as our outcome measure of global cognitive performance. This approach was selected because the BACS composite provides a broad, reliable index of the generalized cognitive impairment characteristic of schizophrenia (20), capturing deficits across multiple domains rather than a single process.

#### Total Cognition Score

For the HCP-EP dataset (Dataset 2) Global cognitive performance was evaluated using the NIH Toolbox Cognition Battery, the standardized neurocognitive assessment protocol utilized by the HCP-EP. The NIH Toolbox is a comprehensive, computerized suite of tests designed to measure core domains of neurological and behavioral function. We specifically focused on the Total Cognition Composite Score. This overarching metric is calculated by aggregating performances across two distinct sub-dimensions: Fluid Cognition (which captures dynamic processing abilities such as executive functioning, episodic memory, processing speed, and working memory) and Crystallized Cognition (which captures accumulated knowledge through vocabulary and reading tasks). Utilizing this composite score provides a single robust index of global cognitive capacity, allowing us to accurately capture the generalized neurocognitive deficits that frequently characterize the early stages of psychotic illness.

### Medication Dose Standardization

Medication dose was measured using Risperidone-equivalents in dataset 1 and Chlorpromazine-equivalents in dataset 2. To harmonize the two scores we followed the empirical data procedure by Leucht et al. (38) and multiplied the Risperidone-equivalent scores from dataset 1 with 100 to get CPZ-equivalent scores.

### Theoretical Identifiability of Attribution Maps

In this section we outline the theoretical underpinnings of the xCEBRA-Time algorithm: we start from the necessary definitions and assumptions for the data generating process and proceed to present the core result for attribution identifiability.

**Definition 1** (Data generating process). We assume that data is generated from a set of latent factors **z**_1_ ∈ R*^d^*^1^ *, …,* **z***_G_* ∈ R*^dG^*. For brevity, the vector **z** ∈ Z ⊆ R*^d^* denotes the concatenation of all factors, and *d* =Σ*_i_ d_i_*. We further assume that the timestep *t* distribution defining the latent process factorizes to:

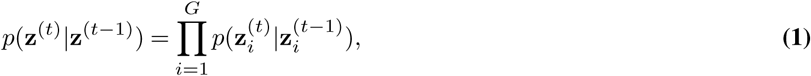

*i.e.*, factors are conditionally independent given their value at the previous time step **z**^(^*^t−^*^1)^. The support of the resulting marginal distributions *p*(**z***_i_*) is assumed to be a convex body or the hypersphere embedded in R*^di^*. The conditional distribution is assumed to take an exponential form:

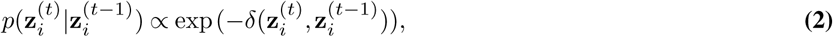

for each latent factor, where *δ* : Z ×Z ’→ R is the negative dot-product or more generally a semi-metric. We observe the process via an injective mixing function **g** : Z ’→ X that maps latent factors to observations, with X ⊆ R*^D^*:

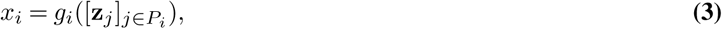

*P_i_* is an index set, and *j* ∈ *P_i_* implies that factor **z***_j_* ∈ R*^dj^* is used to generate the output *x_i_*. Finally, we allow for some factors to be connected to auxiliary variables **c***_i_* through bijective maps *γ_i_* : R*^di^* ’→ R*^di^* s.t. **z***_i_* = *γ_i_*(**c***_i_*).

We proceed with a rigorous definition of identifiability for time-series attribution maps. Under the data-generating framework defined above, consider a feature encoder **f** : X ’→ Z which maps observable data to an manifold space (i.e., the neural-network encoder of the xCEBRA-Time algorithm). The feature encoder is part of a probabilistic model with density *p***_f_**. We define:

**Definition 2** (Subspace Identifiability). Feature encoders **f** *^′^,* **f** *^∗^* : X ’→ Z are identifiable up to subspaces if matching distributions *p***_f_***_′_* = *p***_f_***_∗_* imply that the following equivalence relation holds (the label “*B*” denotes “blockwise”):

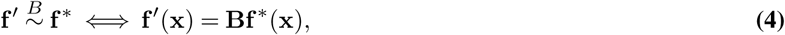

for the block-diagonal matrix **B** ∈ R*^d×d^* with blocks of sizes *d*_1_ × *d*_1_*, …, d_G_* × *d_G_*. In words: two feature encoders are deemed identifiable if their corresponding latent spaces are related by a block-linear transformation that respects the structure of the data-generating process.

Next, we extend the concept of identifiability to attribution maps. An attribution map **A** ∈ A ⊆ R*^D×d^* contains scores *A_ij_* ∈ R. If the *i*-th latent is connected to the *j*-th output, we expect a high score, otherwise a low score. We note here how the notion of attribution score is relative and our definitions below are invariant to scaling and shifting of the scores. For two attribution methods generating attribution maps **A***^′^,* **A***^∗^* ∈ A, we define the following equivalence relation on A:

**Definition 3** (Identifiability of connectivity in attribution maps). Let **A***^′^,* **A***^∗^* ∈ A be attribution maps for the feature encoders

**f** *^′^,* **f** *^∗^*. Let ∼*^C^* be a pairwise relation on A defined as:

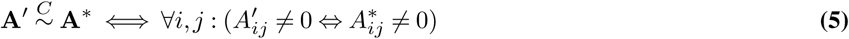

An attribution method is identifiable if the following relation holds (the label “*C*” denotes “connectivity”):

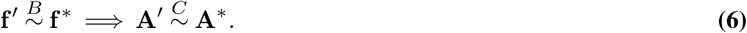

The relation ∼*^C^* equates two different attribution maps **A***^′^* and **A***^∗^* if the locations of their respective “zero entries” match, *i.e.* if both attributions concord in classifying some input as non-relevant. The definition above then states that we consider an attribution method identifiable if, given two equivalent (under the relation ∼*^B^*) encoders, their corresponding attributions are also equivalent (under the relation ∼*^C^*). We stress here how, for scientific discovery and applications to medical settings, obtaining this property is already of high value: it guarantees that the inferred set of relevant relations between data and latent factors (i.e. which part of the observable data are “connected” to the latent space) is reliable and does not depend on either model or training details.

We now have all the necessary definitions to establish a ground-truth attribution map for the mixing function **g**, against which we can later compare the attribution map derived from xCEBRA-Time. Specifically, we are interested in how the factors **z** are connected to the generated data **x** by means of any non-linear mapping. The connectivity defined in Eq. 3 can be read out by considering the Jacobian matrix of **g**, which lets us define the ground truth attribution map as follows:

**Definition 4** (Ground truth attribution map of the mixing function). The ground-truth attribution map **A_g_** ∈ A of the mixing function **g** is defined via the following relationship to the Jacobian matrix **J_g_**:

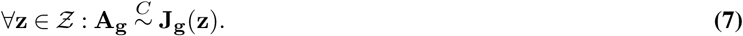

Intuitively, zero-valued derivatives of the observable data with respect to a latent defines non-connectivity. This definition of the “ground-truth map” is intentionally quite flexible and does not imply a particular form of the ground-truth map **A_g_** besides the locations of zeros.

Our attribution map **A** is a *D* × *d*-dimensional matrix and its entry *A_ij_* denotes if the latent at dimension *j* is related to input dimension *i*. We can compute such a map for every time point in the dataset or aggregate multiple time points into a global map. After training **f** using our regularized contrastive learning method, we obtain attribution maps by computing the Jacobian matrix **J_f_** (**x**). We then consider its pseudo-inverse **J**^+^(**x**) at every timestep, which we call the “inverted neuron gradient”. The estimation coincides with the “neuron gradient” attribution method (39), however this has not been paired with identifiable regularized contrastive learning with time-only variables, as proposed here.

Our work focuses on the problem of clearly delineating the *binary* relationship between latents and input data. For this, we threshold the attribution map with a variable decision threshold *ɛ*, **A**^^^ (**x**) := **1**{|**J**^+^(**x**)| *> ɛ*}.

To obtain a global attribution map from local attribution maps, we additionally improve the signal-to-noise ratio by averaging multiple maps. In practice, we found that the operation:

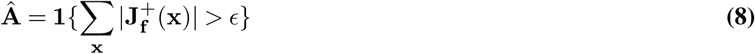

yields even better performance, which we used for all experiments. An alternative, which we considered but did not further explore due to seeing considerably worse performance, is to leverage a max operation instead of the sum. Taking the median is possible, and performs roughly on par with the mean.

We can now finally state the core theoretical identifiability result for the attribution method used by xCEBRA-Time.

**Theorem 1** (Identifiable Attribution Maps). Assume that the data-generating process follows Def. Eq. (1) with mixing function **g** having the associated ground-truth attribution map **A_g_** defined according to Def. Eq. (4). Consider a feature encoder **f** := [**f**_1_; … ; **f***_G_*], with **f***_i_* : X ’→ R*^di^* and assume that it minimizes the Regularized Contrastive Learning loss Eq. (12) on the support *p*(**z**). Then, in the limit of infinite samples *N* → ∞, we have:

1. xCEBRA identifies the latent subspaces of the ground-truth process, i.e. **g**(**f** (**x**)) = **Bx**, with block-diagonal matrix **B**.
2. The Jacobian-pseudo inverse matrix of the xCEBRA feature encoder **J**^+^ identifies the ground-truth attribution map **A_g_**, i.e. for every **x** it holds:

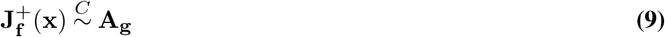

For a complete proof of Theorem 1 we refer to Appendix A of (17).

### Regularized Contrastive Learning

In this section, we present the base adopted self-supervised learning framework employed by the xCEBRA algorithm (16, 17). In the following, we call *p*(·|·) the positive and *q*(·|·) the negative sample distribution. We call (**x**, **x**^+^) a positive pair, and all (**x**, **x***^−^*) for *i* ∈ [*N* ] negative pairs. We call **x** the input time-series data, for example neural activity recorded from the brain or voxel activation in an rs-fMRI.

We define a feature encoder **f** := [**f**_1_; *…* ; **f***_G_*], with **f***_i_* : X ’→ R*^di^* that maps samples into a manifold space partitioned into *G* groups. In practice, we parameterize **f** as a single neural network and only split the last layer into *G* different parts. For training, we apply similarity metrics *ϕ_i_* : R*^di^* ×R*^di^* ’→ R to the different parts of this feature encoder, abbreviated as *ψ_i_*(**x**, **y**) := *ϕ_i_*(**f** (**x**), **f** (**y**)). We then leverage the generalized InfoNCE loss (16),

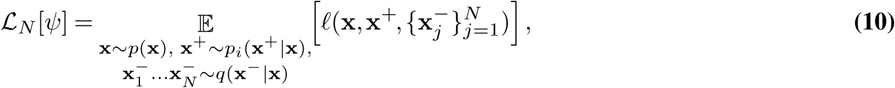

where we have used the InfoNCE loss function:

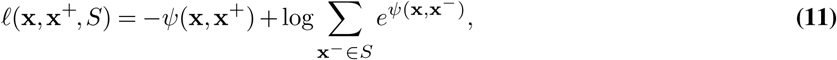

where *S* denotes a set of negative examples. In addition, we regularize the Jacobian matrix of the feature encoder by minimizing its Frobenius norm ∥, · ∥*_F_* (40). With these constraints, we the final objective function, which we refer to as *Regularized Contrastive Learning*, takes the form:

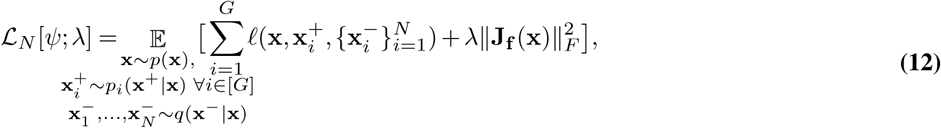

where **J_f_** (**x**) is the Jacobian of the feature encoder **f** optimized as part of *ψ*, ∥, · ∥*_F_* denotes the Frobenius norm and *λ* is a hyperparameter tuned based on the learning dynamics. In practice, *λ* is set to the highest value possible that still allows the InfoNCE component of the loss to stay at its minimum.

### xCEBRA-Time Training

To leverage the shared underlying neural dynamics across cohorts, we concatenated the rs-fMRI time series data from all subjects in both datasets. This unified dataset was used for time-contrastive self-supervised training, without access to any labels. As a feature encoder **f**, we consider a convolutional neural network *h* : F → R*^d^* that maps the rs-fMRI signals F ⊂ R*^D^*, *D > d* into latent vectors. For training we introduce a similarity metric *ϕ* : R*^d^* × R*^d^* ’→ R which is applied to the latent manifolds produced by the encoder as *ψ* (*x, y*) := *ϕ* (*h*(*x*)*, h*(*y*)) */τ*, where *x, y* ∈ F are elements of the training set, *ϕ* is the cosine similarity metric and *τ* is a temperature parameter. Finally, we optimize encoder parameters to minimize the regularized InfoNCE loss Eq. (12) via gradient descent.

In practice, training drives the system to reduce the difference (measured via the similarity metric *ϕ*) between the reference and the positive samples in the manifold space, while simultaneously maximizing the difference between the reference and the negative samples. Reference samples were drawn uniformly across the training set. Positive samples were drawn 20 time-points away from the reference in both temporal directions. Negative samples were uniformly drawn across the training set.

As encoder *h*, we trained a convolutional neural architecture, which utilizes a cascade of 1D convolutions followed by pointwise non-linear activation functions to process a temporal signal. The network consisted of 3 blocks and a final normalization layer. Each block was composed of a convolutional layer with 1024 hidden units (channels) followed by a GELU activation function, with a residual skip connection in the intermediate block. The total receptive-field size of the model was 5 time-steps (offset5model). This configuration resulted in a total capacity of approximately 3.5 million trainable parameters. The model ultimately projected the rs-fMRI inputs into a 32-dimensional latent space. The dimensionality of the latent space was determined by optimizing decoding performance on dataset 2 and was subsequently held constant during model training on both datasets.. Training was performed using a temperature of *τ* = 0.1 for 20, 000 iterations using the Adam optimizer, a learning rate of *η* = 3 × 10*^−^*^4^, and a batch size of 512.

### Regularization

To determine the correct Jacobian regularizer weight *λ* to be able to compute identifiable attribution maps we performed a Jacobian regularizer logarithmic weight sweep over multiple values *λ* ∈ {0, 1, 5, 10, 25, 100, 1000, 10000}. For each candidate weight, we trained three independent xCEBRA models to account for initialization variability. A linear ramp-up scheduler was used for the regularizer, starting at step 5, 000 and reaching the full weight at step 10, 000. We calculated the mean contrastive loss over the final 10 training steps averaged across the three seeds for each weight. Following (17), we selected the highest regularization weight that did not result in a substantial increase in contrastive loss.

### Manifold Alignment and Ensembling

To ensure the robustness of the latent representations and control for variability from network initialization, we constructed a consensus manifold by ensembling multiple independent training runs. We selected three models trained with the optimal Jacobian regularization weight (*λ* = 10) Since unsupervised contrastive learning yields manifolds that are invariant to rotations, the latent spaces from independent runs are not immediately comparable. To resolve this, we first aligned the manifold spaces into a common geometric frame using Orthogonal Procrustes alignments. The resulting aligned manifolds were averaged to get the final manifold.

### Decoding and Evaluation Pipeline

To evaluate the clinical relevance of the learned manifold spaces, we implemented a nested cross-validation decoding pipeline designed to predict clinical, cognitive and demographic scores (e.g., PANSS scores, cognitive batteries) from the manifolds of the xCEBRA-Time models. This procedure was used for hyperparameter tuning and unbiased performance estimation. To prevent data leakage, all data splits were subject-wise, ensuring that frames from the same subject never appeared simultaneously in the training and evaluation sets.

1. *Outer Loop (Performance Estimation):* We used a 10-fold subject-level stratified cross-validation. Subjects were stratified based on their target value to ensure that the distribution of clinical scores was balanced across training and test folds.
2. *Inner Loop (Model Selection):* Within each outer training fold, we performed a further 5-fold subject-level stratified cross-validation. This inner loop was used for hyperparameter optimization.

### Decoder Architecture and Hyperparameters

We utilized a k-Nearest Neighbors (kNN) regressor. We performed a grid search over the following hyperparameter space during the inner CV loop:

- *Number of neighbors (k):* 1, 10, 25, 50, 100, 500, 1000, 5000
- *Weighting function:* Uniform weights, Distance weights.

### Model Selection and Aggregation

Predictions were generated at the frame level. For model selection and final evaluation the frame level predictions were aggregated by meaning for each subject and session in Dataset 1 and each subject for Dataset 2 (as all rs-fMRI sessions were on the same day). The optimal hyperparameter combination was selected based on the Pearson correlation coefficient calculated between the ground truth and the aggregated predictions in the inner validation folds.

### Statistical Evaluation

Once the optimal hyperparameters were identified in the inner loop, a decoder was retrained on the entire outer training split and evaluated on the held-out outer test subjects. We report Pearson correlation and *R*^2^ values. To evaluate possible confounding factors we additionally performed partial correlations using movement and medication scores.

### Jacobian Feature Attribution

To interpret the learned manifold spaces and identify the brain regions driving the latent representations, we performed an inverse Jacobian analysis.

### Computation of Attribution Maps

For three trained xCEBRA-Time models (*λ* = 10), we computed the inverse Jacobian matrix using Singular Value Decomposition. Before the inversion the absolute values of the Jacobian matrix (magnitude of influence) were taken. To obtain a global measure of regional importance, the time-resolved Jacobian matrices were averaged across all time points in the dataset. This resulted in a single summary matrix for each model (Input dimensions x Output dimensions). To correct for initialization inconsistency the attribution maps of the three CEBRA models were Procrustes aligned. As xCEBRA can only discern between zero and non-zero entries we applied a binarization at z = 0 to the aligned attribution maps. The three binarized attribution maps were then averaged for the consensus attribution map.

To visualize the results, the consensus attribution map was then averaged over the output dimensions to get one value of importance for every ROI (ranging from 0 to 1). This value was plotted onto a standard MNI atlas.

### Specificity Analysis

To determine whether the identified features were driven by patient-specific neural dynamics, we trained separate xCEBRA models using only data from the patient cohort or only data from the healthy control cohort. Consensus attribution maps were generated for each group following the same alignment and binarization procedure described above. To analyze group differences, we computed a subtraction attribution map, by mean-centering both attribution maps and then performing an element-wise subtraction of the control consensus map from the patient consensus map.

### Robustness Analysis

To evaluate the stability of the feature attribution results, we conducted a systematic robustness check. We repeated the full training and attribution pipeline while varying one model hyperparameter at a time. We tested the parameters: Number of hidden units, Temperature (*τ*), Time offset (Δ*t*), Jacobian regularization weight (*λ*). The resulting attribution maps were compared to the baseline consensus map. We quantified the similarity between maps using the *R*^2^-value to assess the consistency of the identified regions across different model configurations.

## Supplementary Figures

**Figure S1.**
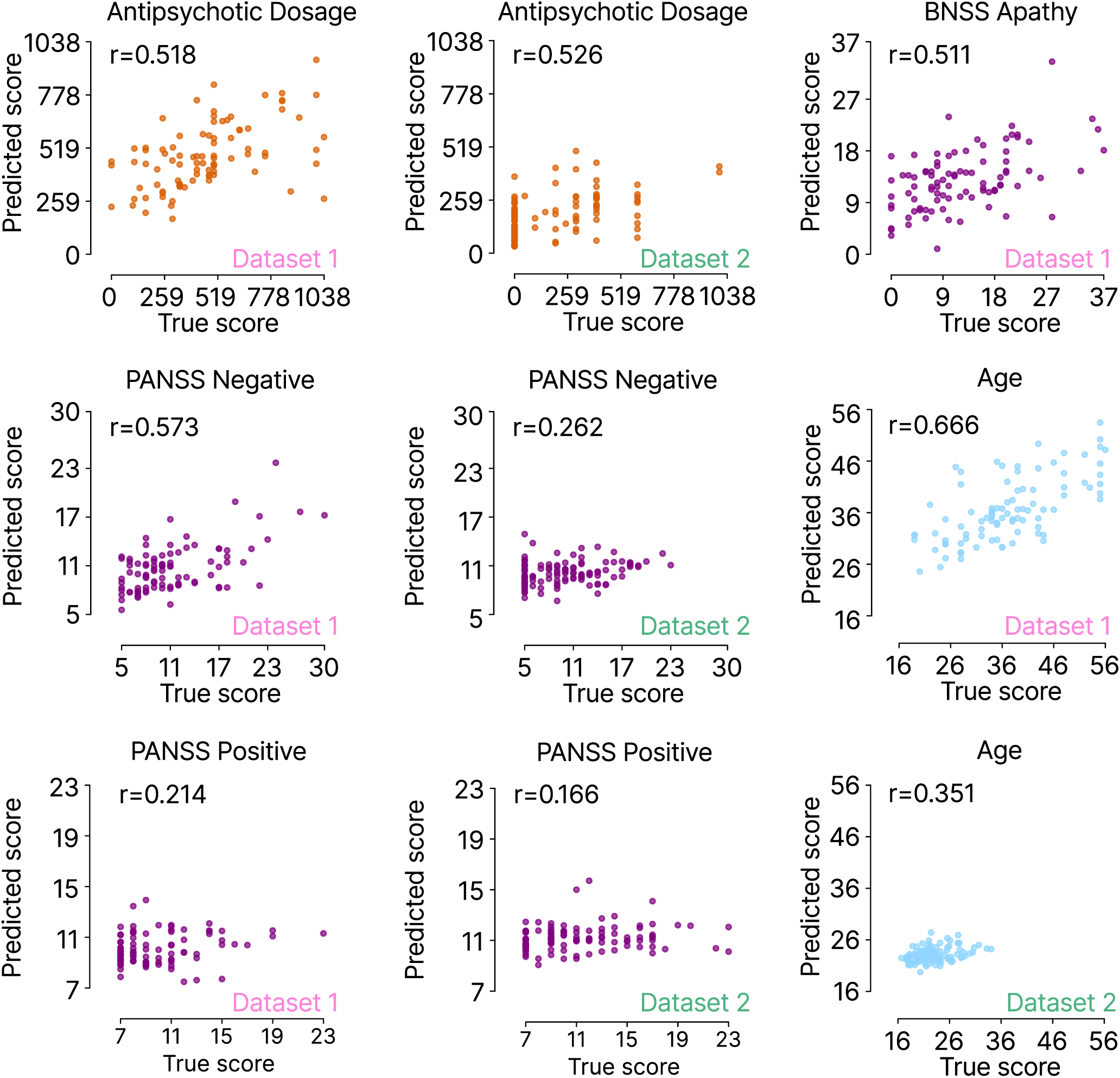
Dataset-specific decoding of disease dimensions using k-NN regression. To ensure that the k-NN predictions reported in Figure 2 were not driven by underlying differences between the two datasets, we independently evaluated all previously performed predictions on a single-dataset level. As with the combined datasets, we report the Pearson correlation between predicted and true scores (r). Additionally, we assessed predictive performance for two extra variables: participant age and an alternative measure of negative symptoms (BNSS-Apathy).

**Figure S2.**
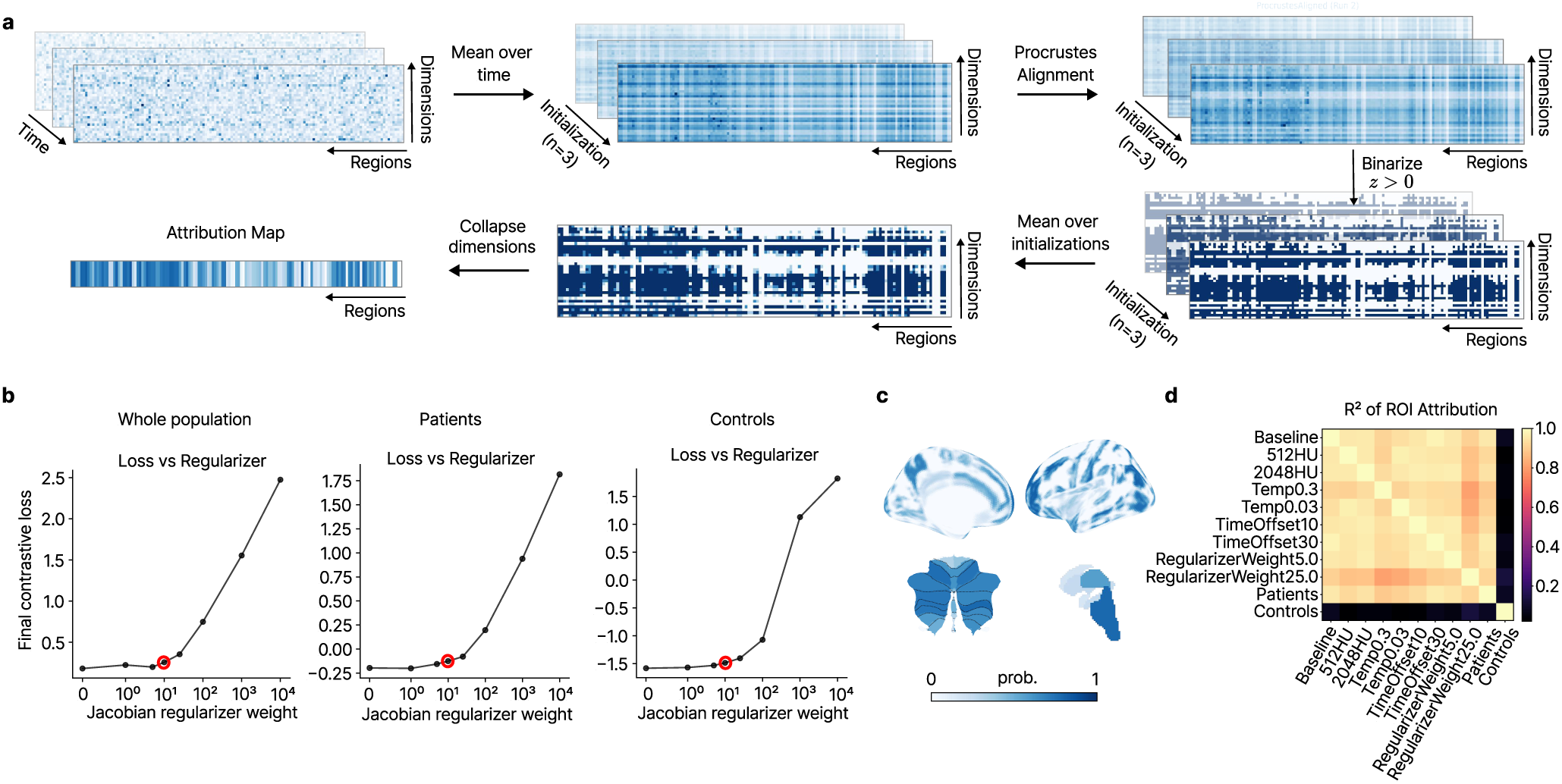
Robustness analysis of attribution map and Jacobian regularizer sweep. a: The influence of various Jacobian regularization weights on the final InfoNCE loss was evaluated to identify the optimal weighting for the model. Loss values represent the mean of the final 10 training steps across three independent initializations of the xCEBRA-Time model. Following theoretical considerations, we selected λ = 10 as the optimal weight, as it was the highest value that did not substantially increase the final training loss. We repeated this process for the xCEBRA-Time models trained on the whole population (left), only on patients (middle) and only on controls (right) b: Jacobian regularizer sweep for the models trained exclusively on the Patient or Healthy Control cohort, to evaluate optimal regularizer weight. c: Reconstructed attribution map from an xCEBRA model trained on the whole population (Patients and Healthy Control). d: Correlation matrix of explained variance (R2), computed while systematically varying one model parameter (Hidden units (HU), Temperature (Temp), Time offset of positive samples (TimeOffset), Jacobian regularizer weight (RegularizerWeight)) or the training dataset (Whole population (Baseline), only patient data (Patients) or only healthy control data (Controls)) at a time.

**Table S1.** Model predictions vs. True scores: Standard and Confounder-Controlled Partial Correlations.

| Target Variable | Standard $r$ | Standard $p$ | Partial $r$<br>(CPZ) | Partial $p$<br>(CPZ) | Partial $r$<br>(Mean Framewise<br>Displacement [mm]) | Partial $p$<br>(Mean Framewise<br>Displacement [mm]) |
| --- | --- | --- | --- | --- | --- | --- |
| PANSS negative | 0.460 | <0.001 | 0.434 | <0.001 | 0.459 | <0.001 |
| CPZ equivalents | 0.676 | <0.001 | - | - | 0.635 | <0.001 |
| BACS Total | 0.496 | <0.001 | 0.498 | <0.001 | 0.496 | <0.001 |
| PANSS positive | 0.241 | <0.001 | 0.240 | <0.001 | 0.241 | <0.001 |
| Age | 0.840 | <0.001 | 0.807 | <0.001 | 0.761 | <0.001 |
| BNSS Apathy | 0.511 | <0.001 | 0.502 | <0.001 | 0.523 | <0.001 |
| Total cognition | 0.509 | <0.001 | 0.413 | <0.001 | 0.496 | <0.001 |

**Table S2.** Regional attributions of the Consensus Map: Whole dataset.

| Category | Region Name | ID | Value |
| --- | --- | --- | --- |
| cortical | Right Middle Temporal Gyrus temporooccipital part | 26 | 0.8438 |
| cortical | Left Middle Frontal Gyrus | 7 | 0.8229 |
| cortical | Left Frontal Pole | 1 | 0.8125 |
| cortical | Left Postcentral Gyrus | 33 | 0.8125 |
| cortical | Right Inferior Temporal Gyrus posterior division | 30 | 0.8125 |
| cortical | Right Frontal Pole | 2 | 0.8125 |
| cortical | Left Superior Frontal Gyrus | 5 | 0.8021 |
| cortical | Right Superior Temporal Gyrus posterior division | 20 | 0.8021 |
| cortical | Left Lateral Occipital Cortex superior division | 43 | 0.8021 |
| cortical | Right Frontal Medial Cortex | 50 | 0.7917 |
| cortical | Right Superior Frontal Gyrus | 6 | 0.7917 |
| cortical | Right Middle Temporal Gyrus posterior division | 24 | 0.7917 |
| cortical | Right Lateral Occipital Cortex superior division | 44 | 0.7917 |
| cortical | Left Middle Temporal Gyrus posterior division | 23 | 0.7812 |
| subcortical | Brain-Stem | 5 | 0.7812 |
| cerebellar | Right_VIIb | 16 | 0.7812 |
| cortical | Right Precentral Gyrus | 14 | 0.7708 |
| cerebellar | Left_VI | 5 | 0.7604 |
| cortical | Right Postcentral Gyrus | 34 | 0.7604 |
| cortical | Left Lateral Occipital Cortex inferior division | 45 | 0.7604 |
| cortical | Right Lateral Occipital Cortex inferior division | 46 | 0.7604 |
| cortical | Right Supramarginal Gyrus posterior division | 40 | 0.7500 |
| cortical | Right Angular Gyrus | 42 | 0.7396 |
| cortical | Left Angular Gyrus | 41 | 0.7396 |
| cerebellar | Left_VIIb | 14 | 0.7396 |
| cerebellar | Right_VI | 7 | 0.7396 |
| cerebellar | Left_CrusII | 11 | 0.7292 |
| cerebellar | Left_VIIIa | 17 | 0.7292 |
| cortical | Right Superior Parietal Lobule | 36 | 0.7292 |
| cortical | Left Middle Temporal Gyrus temporooccipital part | 25 | 0.7292 |
| cortical | Right Frontal Orbital Cortex | 66 | 0.7292 |
| cortical | Left Superior Parietal Lobule | 35 | 0.7188 |
| cortical | Left Frontal Orbital Cortex | 65 | 0.7188 |
| cortical | Right Occipital Pole | 96 | 0.7188 |
| cortical | Right Middle Frontal Gyrus | 8 | 0.7188 |
| cortical | Right Temporal Pole | 16 | 0.7083 |
| cortical | Left Inferior Temporal Gyrus posterior division | 29 | 0.7083 |
| cortical | Left Inferior Frontal Gyrus pars opercularis | 11 | 0.7083 |
| cortical | Left Supramarginal Gyrus posterior division | 39 | 0.7083 |
| cortical | Left Precuneous Cortex | 61 | 0.6979 |
| cortical | Left Temporal Pole | 15 | 0.6979 |
| cerebellar | Left_CrusI | 8 | 0.6979 |
| cortical | Left Precentral Gyrus | 13 | 0.6875 |
| cortical | Left Frontal Medial Cortex | 49 | 0.6875 |
| cerebellar | Right_CrusI | 10 | 0.6771 |
| cortical | Right Paracingulate Gyrus | 56 | 0.6667 |
| cerebellar | Right_CrusII | 13 | 0.6667 |
| cortical | Left Paracingulate Gyrus | 55 | 0.6562 |
| cortical | Left Superior Temporal Gyrus posterior division | 19 | 0.6458 |
| cerebellar | Right_VIIIa | 19 | 0.6458 |
| cortical | Right Inferior Temporal Gyrus temporooccipital part | 32 | 0.6458 |
| cortical | Left Occipital Pole | 95 | 0.6458 |
| subcortical | Right Thalamus | 9 | 0.6354 |

Table S2 – continued from previous page
| Category | Region Name | ID | Value |
| --- | --- | --- | --- |
| cerebellar | Right_IX | 25 | 0.6354 |
| cerebellar | Right_VIIIb | 22 | 0.6354 |
| cerebellar | Right_V | 4 | 0.6146 |
| cerebellar | Left_V | 3 | 0.6146 |
| cerebellar | Vermis_VI | 6 | 0.6146 |
| cortical | Right Precuneous Cortex | 62 | 0.6146 |
| cortical | Right Inferior Frontal Gyrus pars opercularis | 12 | 0.6146 |
| cerebellar | Left_Dentate | 29 | 0.6042 |
| cortical | Right Supramarginal Gyrus anterior division | 38 | 0.6042 |
| cortical | Left Supramarginal Gyrus anterior division | 37 | 0.6042 |
| cerebellar | Right_Dentate | 30 | 0.6042 |
| cortical | Left Cingulate Gyrus anterior division | 57 | 0.5833 |
| cerebellar | Left_VIIIb | 20 | 0.5833 |
| cortical | Left Inferior Frontal Gyrus pars triangularis | 9 | 0.5729 |
| cortical | Left Intracalcarine Cortex | 47 | 0.5729 |
| cortical | Right Central Opercular Cortex | 84 | 0.5729 |
| subcortical | Left Thalamus | 1 | 0.5625 |
| cortical | Right Lingual Gyrus | 72 | 0.5417 |
| cortical | Left Planum Polare | 87 | 0.5312 |
| cortical | Left Lingual Gyrus | 71 | 0.5208 |
| cortical | Left Cuneal Cortex | 63 | 0.5104 |
| cortical | Right Cingulate Gyrus anterior division | 58 | 0.5104 |
| cortical | Left Insular Cortex | 3 | 0.5104 |
| cortical | Left Central Opercular Cortex | 83 | 0.5000 |
| cortical | Left Juxtapositional Lobule Cortex (formerly Supplementary Motor Cortex) | 51 | 0.5000 |
| cerebellar | Left_IX | 23 | 0.5000 |
| cerebellar | Vermis_VIIIa | 18 | 0.4896 |
| cortical | Left Inferior Temporal Gyrus temporooccipital part | 31 | 0.4896 |
| cortical | Left Middle Temporal Gyrus anterior division | 21 | 0.4792 |
| cortical | Right Insular Cortex | 4 | 0.4688 |
| cortical | Right Inferior Frontal Gyrus pars triangularis | 10 | 0.4375 |
| cerebellar | Vermis_CrusII | 12 | 0.4375 |
| cortical | Right Planum Polare | 88 | 0.4271 |
| cortical | Left Frontal Operculum Cortex | 81 | 0.4271 |
| cortical | Right Parietal Operculum Cortex | 86 | 0.4167 |
| cortical | Left Parietal Operculum Cortex | 85 | 0.4167 |
| cortical | Right Juxtapositional Lobule Cortex (formerly Supplementary Motor Cortex) | 52 | 0.4062 |
| cortical | Right Occipital Fusiform Gyrus | 80 | 0.3958 |
| cerebellar | Right_I_IV | 2 | 0.3958 |
| cortical | Left Temporal Fusiform Cortex posterior division | 75 | 0.3854 |
| cortical | Right Cuneal Cortex | 64 | 0.3750 |
| cortical | Left Occipital Fusiform Gyrus | 79 | 0.3646 |
| cortical | Right Temporal Occipital Fusiform Cortex | 78 | 0.3542 |
| cortical | Right Cingulate Gyrus posterior division | 60 | 0.3438 |
| subcortical | Right Hippocampus | 13 | 0.3333 |
| cortical | Right Intracalcarine Cortex | 48 | 0.3333 |
| cortical | Right Middle Temporal Gyrus anterior division | 22 | 0.3333 |
| cortical | Right Planum Temporale | 92 | 0.3229 |
| cortical | Right Temporal Fusiform Cortex posterior division | 76 | 0.3125 |
| cerebellar | Left_I_IV | 1 | 0.3125 |
| cerebellar | Vermis_VIIIb | 21 | 0.3021 |
| cortical | Left Planum Temporale | 91 | 0.2917 |
| cortical | Left Cingulate Gyrus posterior division | 59 | 0.2917 |
| cortical | Left Inferior Temporal Gyrus anterior division | 27 | 0.2812 |

**Table S2 – continued from previous page**
| **Category** | **Region Name** | **ID** | **Value** |
| --- | --- | --- | --- |
| cortical | Left Temporal Occipital Fusiform Cortex | 77 | 0.2708 |
| cortical | Left Superior Temporal Gyrus anterior division | 17 | 0.2604 |
| cortical | Right Subcallosal Cortex | 54 | 0.2500 |
| cortical | Right Superior Temporal Gyrus anterior division | 18 | 0.2500 |
| subcortical | Left Putamen | 3 | 0.2500 |
| cortical | Right Parahippocampal Gyrus anterior division | 68 | 0.2396 |
| subcortical | Left Hippocampus | 6 | 0.2396 |
| subcortical | Right Caudate | 10 | 0.2083 |
| cortical | Right Frontal Operculum Cortex | 82 | 0.2083 |
| cerebellar | Vermis_IX | 24 | 0.1979 |
| subcortical | Right Putamen | 11 | 0.1979 |
| cortical | Left Heschl's Gyrus (includes H1 and H2) | 89 | 0.1979 |
| cortical | Right Heschl's Gyrus (includes H1 and H2) | 90 | 0.1875 |
| cortical | Left Subcallosal Cortex | 53 | 0.1771 |
| subcortical | Left Caudate | 2 | 0.1771 |
| cortical | Left Parahippocampal Gyrus anterior division | 67 | 0.1667 |
| cortical | Right Inferior Temporal Gyrus anterior division | 28 | 0.1667 |
| subcortical | Left Amygdala | 7 | 0.1562 |
| subcortical | Right Amygdala | 14 | 0.1458 |
| subcortical | Right Pallidum | 12 | 0.1354 |
| subcortical | Left Pallidum | 4 | 0.1146 |
| cortical | Left Temporal Fusiform Cortex anterior division | 73 | 0.0625 |
| cortical | Left Parahippocampal Gyrus posterior division | 69 | 0.0521 |
| cortical | Right Parahippocampal Gyrus posterior division | 70 | 0.0417 |
| cerebellar | Vermis_VIIb | 15 | 0.0312 |
| cerebellar | Vermis_CrusI | 9 | 0.0312 |
| cerebellar | Right_X | 28 | 0.0104 |
| cerebellar | Left_X | 26 | 0.0000 |
| cerebellar | Vermis_X | 27 | 0.0000 |
| subcortical | Left Accumbens | 8 | 0.0000 |
| subcortical | Right Accumbens | 15 | 0.0000 |
| cortical | Right Temporal Fusiform Cortex anterior division | 74 | 0.0000 |
| cerebellar | Left_Interposed | 31 | 0.0000 |
| cerebellar | Right_Interposed | 32 | 0.0000 |

**Table S3.** Regional attributions of the Consensus Map: Only Patients.

| **Category** | **Region Name** | **ID** | **Value** |
| --- | --- | --- | --- |
| cortical | Left Lateral Occipital Cortex superior division | 43 | 0.8333 |
| cortical | Left Middle Frontal Gyrus | 7 | 0.8021 |
| cortical | Left Middle Temporal Gyrus temporooccipital part | 25 | 0.7812 |
| cortical | Right Middle Temporal Gyrus temporooccipital part | 26 | 0.7708 |
| cortical | Right Lateral Occipital Cortex superior division | 44 | 0.7604 |
| subcortical | Brain-Stem | 5 | 0.7604 |
| cortical | Right Precentral Gyrus | 14 | 0.7604 |
| cortical | Right Superior Frontal Gyrus | 6 | 0.7604 |
| cortical | Left Angular Gyrus | 41 | 0.7604 |
| cortical | Left Supramarginal Gyrus posterior division | 39 | 0.7604 |
| cortical | Right Frontal Pole | 2 | 0.7500 |
| cortical | Right Angular Gyrus | 42 | 0.7396 |
| cortical | Right Supramarginal Gyrus posterior division | 40 | 0.7396 |
| cortical | Right Postcentral Gyrus | 34 | 0.7396 |

Table S3 – continued from previous page
| Category | Region Name | ID | Value |
| --- | --- | --- | --- |
| cortical | Right Frontal Medial Cortex | 50 | 0.7396 |
| cortical | Left Postcentral Gyrus | 33 | 0.7396 |
| cortical | Left Superior Frontal Gyrus | 5 | 0.7292 |
| cerebellar | Right_VIIb | 16 | 0.7292 |
| cortical | Left Frontal Pole | 1 | 0.7188 |
| cortical | Right Middle Temporal Gyrus posterior division | 24 | 0.7188 |
| cerebellar | Left_VIIb | 14 | 0.7083 |
| cortical | Right Superior Temporal Gyrus posterior division | 20 | 0.7083 |
| cortical | Left Lateral Occipital Cortex inferior division | 45 | 0.6979 |
| cortical | Right Inferior Temporal Gyrus posterior division | 30 | 0.6979 |
| cerebellar | Left_CrusII | 11 | 0.6875 |
| cerebellar | Left_VI | 5 | 0.6875 |
| cortical | Right Temporal Pole | 16 | 0.6771 |
| cerebellar | Left_VIIIa | 17 | 0.6771 |
| cortical | Right Lateral Occipital Cortex inferior division | 46 | 0.6667 |
| cerebellar | Right_VI | 7 | 0.6562 |
| cortical | Left Middle Temporal Gyrus posterior division | 23 | 0.6562 |
| cortical | Left Temporal Pole | 15 | 0.6562 |
| cortical | Right Superior Parietal Lobule | 36 | 0.6562 |
| cortical | Right Frontal Orbital Cortex | 66 | 0.6562 |
| cortical | Left Superior Parietal Lobule | 35 | 0.6562 |
| cortical | Right Middle Frontal Gyrus | 8 | 0.6458 |
| cortical | Left Superior Temporal Gyrus posterior division | 19 | 0.6458 |
| cortical | Left Frontal Orbital Cortex | 65 | 0.6354 |
| cortical | Right Occipital Pole | 96 | 0.6354 |
| cerebellar | Right_CrusII | 13 | 0.6354 |
| cortical | Right Inferior Temporal Gyrus temporooccipital part | 32 | 0.6250 |
| cerebellar | Left_CrusI | 8 | 0.6250 |
| cortical | Right Paracingulate Gyrus | 56 | 0.6250 |
| cortical | Left Inferior Frontal Gyrus pars triangularis | 9 | 0.6146 |
| cortical | Left Paracingulate Gyrus | 55 | 0.6146 |
| cortical | Left Inferior Frontal Gyrus pars opercularis | 11 | 0.6042 |
| cortical | Left Frontal Medial Cortex | 49 | 0.5938 |
| cortical | Left Precentral Gyrus | 13 | 0.5938 |
| cortical | Left Inferior Temporal Gyrus posterior division | 29 | 0.5938 |
| cerebellar | Right_CrusI | 10 | 0.5833 |
| cortical | Right Inferior Frontal Gyrus pars opercularis | 12 | 0.5833 |
| cerebellar | Right_VIIIa | 19 | 0.5833 |
| cerebellar | Vermis_VI | 6 | 0.5729 |
| cortical | Left Supramarginal Gyrus anterior division | 37 | 0.5625 |
| cortical | Right Cingulate Gyrus anterior division | 58 | 0.5625 |
| cortical | Right Precuneous Cortex | 62 | 0.5625 |
| cerebellar | Right_Dentate | 30 | 0.5625 |
| cortical | Left Cuneal Cortex | 63 | 0.5521 |
| cortical | Left Occipital Pole | 95 | 0.5521 |
| subcortical | Right Thalamus | 9 | 0.5417 |
| cerebellar | Left_Dentate | 29 | 0.5417 |
| cerebellar | Left_V | 3 | 0.5417 |
| cortical | Right Supramarginal Gyrus anterior division | 38 | 0.5417 |
| cerebellar | Right_VIIIb | 22 | 0.5312 |
| cortical | Left Insular Cortex | 3 | 0.5312 |
| cortical | Left Precuneous Cortex | 61 | 0.5312 |
| subcortical | Left Thalamus | 1 | 0.5312 |
| cerebellar | Left_VIIIb | 20 | 0.5312 |

Table S3 – continued from previous page
| Category | Region Name | ID | Value |
| --- | --- | --- | --- |
| cortical | Left Cingulate Gyrus anterior division | 57 | 0.5208 |
| cerebellar | Right_V | 4 | 0.5208 |
| cortical | Left Juxtapositional Lobule Cortex (formerly Supplementary Motor Cortex) | 51 | 0.5104 |
| cerebellar | Vermis_VIIIa | 18 | 0.5000 |
| cerebellar | Right_IX | 25 | 0.5000 |
| cortical | Left Intracalcarine Cortex | 47 | 0.4896 |
| cerebellar | Right_I_IV | 2 | 0.4792 |
| cortical | Right Lingual Gyrus | 72 | 0.4688 |
| cerebellar | Left_IX | 23 | 0.4688 |
| cortical | Left Planum Polare | 87 | 0.4688 |
| cortical | Right Insular Cortex | 4 | 0.4688 |
| cortical | Right Inferior Frontal Gyrus pars triangularis | 10 | 0.4583 |
| cortical | Right Central Opercular Cortex | 84 | 0.4583 |
| cortical | Right Juxtapositional Lobule Cortex (formerly Supplementary Motor Cortex) | 52 | 0.4583 |
| cortical | Left Lingual Gyrus | 71 | 0.4479 |
| cortical | Left Parietal Operculum Cortex | 85 | 0.4479 |
| cortical | Left Inferior Temporal Gyrus temporooccipital part | 31 | 0.4479 |
| cortical | Right Planum Polare | 88 | 0.4479 |
| cortical | Left Middle Temporal Gyrus anterior division | 21 | 0.4479 |
| cortical | Left Central Opercular Cortex | 83 | 0.4375 |
| cortical | Left Temporal Fusiform Cortex posterior division | 75 | 0.4375 |
| cortical | Right Parietal Operculum Cortex | 86 | 0.4271 |
| cortical | Left Frontal Operculum Cortex | 81 | 0.4271 |
| cortical | Right Cingulate Gyrus posterior division | 60 | 0.4062 |
| cortical | Left Planum Temporale | 91 | 0.4062 |
| cerebellar | Left_I_IV | 1 | 0.4062 |
| cortical | Left Occipital Fusiform Gyrus | 79 | 0.3958 |
| cortical | Right Planum Temporale | 92 | 0.3854 |
| cortical | Right Occipital Fusiform Gyrus | 80 | 0.3750 |
| cortical | Right Temporal Fusiform Cortex posterior division | 76 | 0.3646 |
| cortical | Right Cuneal Cortex | 64 | 0.3542 |
| cortical | Right Temporal Occipital Fusiform Cortex | 78 | 0.3542 |
| cortical | Left Inferior Temporal Gyrus anterior division | 27 | 0.3438 |
| cortical | Left Cingulate Gyrus posterior division | 59 | 0.3438 |
| cortical | Left Temporal Occipital Fusiform Cortex | 77 | 0.3333 |
| subcortical | Right Hippocampus | 13 | 0.3333 |
| subcortical | Left Hippocampus | 6 | 0.3229 |
| cortical | Right Middle Temporal Gyrus anterior division | 22 | 0.3229 |
| subcortical | Left Putamen | 3 | 0.3125 |
| cortical | Right Intracalcarine Cortex | 48 | 0.3021 |
| cortical | Left Superior Temporal Gyrus anterior division | 17 | 0.2917 |
| cerebellar | Vermis_CrusII | 12 | 0.2812 |
| cortical | Right Superior Temporal Gyrus anterior division | 18 | 0.2812 |
| cerebellar | Vermis_VIIIb | 21 | 0.2500 |
| cortical | Right Subcallosal Cortex | 54 | 0.2500 |
| subcortical | Right Putamen | 11 | 0.2396 |
| cortical | Right Inferior Temporal Gyrus anterior division | 28 | 0.2396 |
| subcortical | Right Caudate | 10 | 0.2292 |
| cortical | Right Frontal Operculum Cortex | 82 | 0.2292 |
| cortical | Right Parahippocampal Gyrus anterior division | 68 | 0.2188 |
| cortical | Left Parahippocampal Gyrus anterior division | 67 | 0.2188 |
| cortical | Left Subcallosal Cortex | 53 | 0.2188 |
| subcortical | Left Caudate | 2 | 0.2083 |
| cortical | Right Heschl's Gyrus (includes H1 and H2) | 90 | 0.1979 |

**Table S3 – continued from previous page**
| **Category** | **Region Name** | **ID** | **Value** |
| --- | --- | --- | --- |
| cortical | Left Heschl's Gyrus (includes H1 and H2) | 89 | 0.1979 |
| subcortical | Left Pallidum | 4 | 0.1875 |
| cerebellar | Vermis_IX | 24 | 0.1875 |
| subcortical | Right Pallidum | 12 | 0.1875 |
| subcortical | Left Amygdala | 7 | 0.1875 |
| cortical | Right Parahippocampal Gyrus posterior division | 70 | 0.1458 |
| subcortical | Right Amygdala | 14 | 0.1354 |
| cortical | Left Parahippocampal Gyrus posterior division | 69 | 0.1250 |
| cerebellar | Vermis_VIIb | 15 | 0.0938 |
| cerebellar | Vermis_X | 27 | 0.0625 |
| cerebellar | Right_X | 28 | 0.0625 |
| cortical | Left Temporal Fusiform Cortex anterior division | 73 | 0.0625 |
| cerebellar | Left_X | 26 | 0.0521 |
| cortical | Right Temporal Fusiform Cortex anterior division | 74 | 0.0521 |
| cerebellar | Vermis_CrusI | 9 | 0.0104 |
| cerebellar | Left_Interposed | 31 | 0.0104 |
| subcortical | Left Accumbens | 8 | 0.0000 |
| subcortical | Right Accumbens | 15 | 0.0000 |
| cerebellar | Right_Interposed | 32 | 0.0000 |

**Table S4.** Regional attributions of the Consensus Map: Only Controls.

| **Category** | **Region Name** | **ID** | **Value** |
| --- | --- | --- | --- |
| cortical | Left Planum Polare | 87 | 0.8854 |
| cortical | Right Planum Polare | 88 | 0.7917 |
| cortical | Right Middle Temporal Gyrus temporooccipital part | 26 | 0.7812 |
| cerebellar | Right_IX | 25 | 0.6771 |
| cerebellar | Left_IX | 23 | 0.6667 |
| cortical | Left Supramarginal Gyrus posterior division | 39 | 0.6354 |
| cortical | Left Middle Temporal Gyrus temporooccipital part | 25 | 0.6354 |
| cortical | Right Supramarginal Gyrus posterior division | 40 | 0.6250 |
| cortical | Right Parahippocampal Gyrus anterior division | 68 | 0.6250 |
| cortical | Right Planum Temporale | 92 | 0.6250 |
| cortical | Left Heschl's Gyrus (includes H1 and H2) | 89 | 0.6146 |
| cortical | Right Frontal Medial Cortex | 50 | 0.6042 |
| cortical | Left Angular Gyrus | 41 | 0.6042 |
| cortical | Right Occipital Pole | 96 | 0.5938 |
| cortical | Right Superior Parietal Lobule | 36 | 0.5833 |
| cortical | Left Superior Temporal Gyrus posterior division | 19 | 0.5729 |
| cortical | Right Inferior Temporal Gyrus temporooccipital part | 32 | 0.5729 |
| cerebellar | Vermis_VI | 6 | 0.5625 |
| cortical | Left Inferior Frontal Gyrus pars opercularis | 11 | 0.5625 |
| subcortical | Right Thalamus | 9 | 0.5625 |
| cortical | Right Superior Temporal Gyrus posterior division | 20 | 0.5521 |
| cerebellar | Vermis_CrusII | 12 | 0.5521 |
| cortical | Left Middle Frontal Gyrus | 7 | 0.5417 |
| cortical | Left Frontal Medial Cortex | 49 | 0.5417 |
| cortical | Left Middle Temporal Gyrus posterior division | 23 | 0.5417 |
| cortical | Right Inferior Frontal Gyrus pars triangularis | 10 | 0.5312 |
| cortical | Left Inferior Temporal Gyrus temporooccipital part | 31 | 0.5208 |
| cortical | Right Heschl's Gyrus (includes H1 and H2) | 90 | 0.5208 |
| cortical | Right Cuneal Cortex | 64 | 0.5208 |

Table S4 – continued from previous page
| Category | Region Name | ID | Value |
| --- | --- | --- | --- |
| cortical | Left Superior Parietal Lobule | 35 | 0.5104 |
| cortical | Right Angular Gyrus | 42 | 0.5104 |
| cerebellar | Left_I_IV | 1 | 0.5000 |
| subcortical | Left Thalamus | 1 | 0.5000 |
| cortical | Left Planum Temporale | 91 | 0.5000 |
| cortical | Right Inferior Frontal Gyrus pars opercularis | 12 | 0.4896 |
| cerebellar | Right_Dentate | 30 | 0.4896 |
| cortical | Right Inferior Temporal Gyrus posterior division | 30 | 0.4792 |
| cortical | Right Middle Temporal Gyrus posterior division | 24 | 0.4792 |
| cortical | Left Inferior Temporal Gyrus posterior division | 29 | 0.4792 |
| cortical | Left Frontal Orbital Cortex | 65 | 0.4688 |
| cortical | Right Middle Temporal Gyrus anterior division | 22 | 0.4479 |
| cerebellar | Right_I_IV | 2 | 0.4479 |
| cortical | Right Frontal Orbital Cortex | 66 | 0.4479 |
| cortical | Left Insular Cortex | 3 | 0.4479 |
| cortical | Right Parietal Operculum Cortex | 86 | 0.4375 |
| cortical | Left Inferior Temporal Gyrus anterior division | 27 | 0.4375 |
| cortical | Left Middle Temporal Gyrus anterior division | 21 | 0.4375 |
| cortical | Right Subcallosal Cortex | 54 | 0.4375 |
| cortical | Left Inferior Frontal Gyrus pars triangularis | 9 | 0.4271 |
| subcortical | Brain-Stem | 5 | 0.4167 |
| cortical | Left Parahippocampal Gyrus anterior division | 67 | 0.4167 |
| cortical | Left Temporal Fusiform Cortex posterior division | 75 | 0.4167 |
| cerebellar | Right_V | 4 | 0.4167 |
| cortical | Right Lateral Occipital Cortex superior division | 44 | 0.4167 |
| cerebellar | Right_VIIIb | 22 | 0.4167 |
| subcortical | Left Amygdala | 7 | 0.4062 |
| cortical | Left Parietal Operculum Cortex | 85 | 0.4062 |
| subcortical | Right Hippocampus | 13 | 0.3958 |
| cortical | Left Cuneal Cortex | 63 | 0.3958 |
| cerebellar | Left_Dentate | 29 | 0.3958 |
| cortical | Right Superior Frontal Gyrus | 6 | 0.3854 |
| cortical | Left Frontal Operculum Cortex | 81 | 0.3854 |
| cerebellar | Left_VIIIb | 20 | 0.3854 |
| subcortical | Right Amygdala | 14 | 0.3854 |
| cortical | Right Temporal Fusiform Cortex posterior division | 76 | 0.3750 |
| cortical | Right Middle Frontal Gyrus | 8 | 0.3750 |
| cortical | Right Superior Temporal Gyrus anterior division | 18 | 0.3542 |
| cortical | Right Supramarginal Gyrus anterior division | 38 | 0.3438 |
| cerebellar | Vermis_VIIIb | 21 | 0.3438 |
| subcortical | Left Hippocampus | 6 | 0.3438 |
| cerebellar | Vermis_IX | 24 | 0.3333 |
| cerebellar | Left_V | 3 | 0.3333 |
| cerebellar | Vermis_VIIIa | 18 | 0.3229 |
| cortical | Right Occipital Fusiform Gyrus | 80 | 0.3229 |
| cortical | Right Temporal Pole | 16 | 0.3125 |
| cortical | Right Lateral Occipital Cortex inferior division | 46 | 0.3125 |
| cortical | Left Parahippocampal Gyrus posterior division | 69 | 0.3021 |
| cortical | Left Temporal Occipital Fusiform Cortex | 77 | 0.3021 |
| cortical | Right Cingulate Gyrus anterior division | 58 | 0.2917 |
| cortical | Left Subcallosal Cortex | 53 | 0.2917 |
| cortical | Left Superior Frontal Gyrus | 5 | 0.2917 |
| cortical | Left Supramarginal Gyrus anterior division | 37 | 0.2917 |
| cortical | Left Occipital Pole | 95 | 0.2917 |

Table S4 – continued from previous page
| Category | Region Name | ID | Value |
| --- | --- | --- | --- |
| subcortical | Left Caudate | 2 | 0.2812 |
| cortical | Left Juxtapositional Lobule Cortex (formerly Supplementary Motor Cortex) | 51 | 0.2812 |
| cerebellar | Left_VIIIa | 17 | 0.2812 |
| cortical | Left Paracingulate Gyrus | 55 | 0.2708 |
| cortical | Right Insular Cortex | 4 | 0.2708 |
| subcortical | Right Caudate | 10 | 0.2708 |
| cortical | Right Inferior Temporal Gyrus anterior division | 28 | 0.2708 |
| cortical | Left Superior Temporal Gyrus anterior division | 17 | 0.2708 |
| cortical | Right Lingual Gyrus | 72 | 0.2708 |
| cortical | Left Intracalcarine Cortex | 47 | 0.2604 |
| subcortical | Right Pallidum | 12 | 0.2604 |
| cerebellar | Left_VI | 5 | 0.2604 |
| cortical | Right Frontal Operculum Cortex | 82 | 0.2604 |
| cortical | Left Temporal Fusiform Cortex anterior division | 73 | 0.2604 |
| cortical | Left Cingulate Gyrus anterior division | 57 | 0.2604 |
| cortical | Right Juxtapositional Lobule Cortex (formerly Supplementary Motor Cortex) | 52 | 0.2604 |
| cerebellar | Right_VIIIa | 19 | 0.2604 |
| subcortical | Left Putamen | 3 | 0.2604 |
| cortical | Right Temporal Occipital Fusiform Cortex | 78 | 0.2500 |
| cortical | Right Central Opercular Cortex | 84 | 0.2500 |
| cerebellar | Right_VIIb | 16 | 0.2500 |
| cortical | Right Intracalcarine Cortex | 48 | 0.2500 |
| cortical | Left Temporal Pole | 15 | 0.2500 |
| cortical | Left Occipital Fusiform Gyrus | 79 | 0.2500 |
| cortical | Left Central Opercular Cortex | 83 | 0.2396 |
| cortical | Right Paracingulate Gyrus | 56 | 0.2396 |
| cortical | Left Lingual Gyrus | 71 | 0.2396 |
| subcortical | Left Pallidum | 4 | 0.2396 |
| subcortical | Right Putamen | 11 | 0.2396 |
| cerebellar | Right_VI | 7 | 0.2396 |
| cerebellar | Vermis_X | 27 | 0.2292 |
| cortical | Left Frontal Pole | 1 | 0.2292 |
| cortical | Right Postcentral Gyrus | 34 | 0.2292 |
| cortical | Left Postcentral Gyrus | 33 | 0.2292 |
| cortical | Left Cingulate Gyrus posterior division | 59 | 0.2188 |
| cortical | Left Lateral Occipital Cortex inferior division | 45 | 0.2188 |
| cerebellar | Vermis_VIIb | 15 | 0.2188 |
| cerebellar | Right_CrusI | 10 | 0.2188 |
| cortical | Right Cingulate Gyrus posterior division | 60 | 0.2188 |
| cortical | Left Lateral Occipital Cortex superior division | 43 | 0.2188 |
| cortical | Right Parahippocampal Gyrus posterior division | 70 | 0.2188 |
| cerebellar | Left_VIIb | 14 | 0.2083 |
| cortical | Right Precentral Gyrus | 14 | 0.2083 |
| cerebellar | Left_CrusII | 11 | 0.2083 |
| cortical | Right Precuneous Cortex | 62 | 0.1979 |
| cortical | Left Precentral Gyrus | 13 | 0.1979 |
| cortical | Left Precuneous Cortex | 61 | 0.1875 |
| cerebellar | Left_X | 26 | 0.1771 |
| cerebellar | Right_X | 28 | 0.1771 |
| cerebellar | Left_CrusI | 8 | 0.1562 |
| cerebellar | Vermis_CrusI | 9 | 0.1562 |
| cerebellar | Right_CrusII | 13 | 0.1354 |
| cortical | Right Frontal Pole | 2 | 0.1354 |
| cortical | Right Temporal Fusiform Cortex anterior division | 74 | 0.1354 |

Table S4 – continued from previous page
| Category | Region Name | ID | Value |
| --- | --- | --- | --- |
| cerebellar | Left_Interposed | 31 | 0.1354 |
| cerebellar | Right_Interposed | 32 | 0.1250 |
| subcortical | Right Accumbens | 15 | 0.0938 |
| subcortical | Left Accumbens | 8 | 0.0938 |

**Table S5.** Regional attributions of the Subtraction Map: Patients minus Controls.

| Category | Region Name | ID | Value |
| --- | --- | --- | --- |
| cortical | Left Lateral Occipital Cortex superior division | 43 | 0.5134 |
| cortical | Right Frontal Pole | 2 | 0.5134 |
| cortical | Right Precentral Gyrus | 14 | 0.4509 |
| cortical | Left Postcentral Gyrus | 33 | 0.4093 |
| cortical | Right Postcentral Gyrus | 34 | 0.4093 |
| cerebellar | Left_VIIb | 14 | 0.3989 |
| cerebellar | Right_CrusII | 13 | 0.3989 |
| cortical | Left Frontal Pole | 1 | 0.3884 |
| cortical | Left Lateral Occipital Cortex inferior division | 45 | 0.3780 |
| cerebellar | Right_VIIb | 16 | 0.3780 |
| cerebellar | Left_CrusII | 11 | 0.3780 |
| cerebellar | Left_CrusI | 8 | 0.3676 |
| cortical | Left Superior Frontal Gyrus | 5 | 0.3364 |
| cerebellar | Left_VI | 5 | 0.3259 |
| cerebellar | Right_VI | 7 | 0.3155 |
| cortical | Left Temporal Pole | 15 | 0.3051 |
| cortical | Left Precentral Gyrus | 13 | 0.2947 |
| cerebellar | Left_VIIIa | 17 | 0.2947 |
| cortical | Right Paracingulate Gyrus | 56 | 0.2843 |
| cortical | Right Superior Frontal Gyrus | 6 | 0.2739 |
| cerebellar | Right_CrusI | 10 | 0.2634 |
| cortical | Right Temporal Pole | 16 | 0.2634 |
| cortical | Right Precuneous Cortex | 62 | 0.2634 |
| cortical | Right Lateral Occipital Cortex inferior division | 46 | 0.2530 |
| cortical | Left Precuneous Cortex | 61 | 0.2426 |
| subcortical | Brain-Stem | 5 | 0.2426 |
| cortical | Right Lateral Occipital Cortex superior division | 44 | 0.2426 |
| cortical | Left Paracingulate Gyrus | 55 | 0.2426 |
| cerebellar | Right_VIIIa | 19 | 0.2218 |
| cortical | Right Middle Frontal Gyrus | 8 | 0.1697 |
| cortical | Left Supramarginal Gyrus anterior division | 37 | 0.1697 |
| cortical | Right Cingulate Gyrus anterior division | 58 | 0.1697 |
| cortical | Left Cingulate Gyrus anterior division | 57 | 0.1593 |
| cortical | Left Occipital Pole | 95 | 0.1593 |
| cortical | Left Middle Frontal Gyrus | 7 | 0.1593 |
| cortical | Right Middle Temporal Gyrus posterior division | 24 | 0.1384 |
| cortical | Right Angular Gyrus | 42 | 0.1280 |
| cortical | Left Juxtapositional Lobule Cortex (formerly Supplementary Motor Cortex) | 51 | 0.1280 |
| cortical | Left Intracalcarine Cortex | 47 | 0.1280 |
| cortical | Right Inferior Temporal Gyrus posterior division | 30 | 0.1176 |
| cerebellar | Left_V | 3 | 0.1072 |
| cortical | Left Lingual Gyrus | 71 | 0.1072 |
| cortical | Right Central Opercular Cortex | 84 | 0.1072 |
| cortical | Right Frontal Orbital Cortex | 66 | 0.1072 |

Table S5 – continued from previous page
| Category | Region Name | ID | Value |
| --- | --- | --- | --- |
| cortical | Right Lingual Gyrus | 72 | 0.0968 |
| cortical | Right Juxtapositional Lobule Cortex (formerly Supplementary Motor Cortex) | 52 | 0.0968 |
| cortical | Left Central Opercular Cortex | 83 | 0.0968 |
| cortical | Right Insular Cortex | 4 | 0.0968 |
| cortical | Right Supramarginal Gyrus anterior division | 38 | 0.0968 |
| cortical | Left Inferior Frontal Gyrus pars triangularis | 9 | 0.0864 |
| cortical | Right Cingulate Gyrus posterior division | 60 | 0.0864 |
| cerebellar | Vermis_VIIIa | 18 | 0.0759 |
| cortical | Left Frontal Orbital Cortex | 65 | 0.0655 |
| cortical | Left Cuneal Cortex | 63 | 0.0551 |
| cortical | Right Superior Temporal Gyrus posterior division | 20 | 0.0551 |
| cortical | Left Angular Gyrus | 41 | 0.0551 |
| cortical | Left Occipital Fusiform Gyrus | 79 | 0.0447 |
| cerebellar | Left_Dentate | 29 | 0.0447 |
| cortical | Left Middle Temporal Gyrus temporooccipital part | 25 | 0.0447 |
| cortical | Left Superior Parietal Lobule | 35 | 0.0447 |
| cerebellar | Left_VIIIb | 20 | 0.0447 |
| cortical | Right Frontal Medial Cortex | 50 | 0.0343 |
| cortical | Left Cingulate Gyrus posterior division | 59 | 0.0239 |
| cortical | Left Supramarginal Gyrus posterior division | 39 | 0.0239 |
| cortical | Left Middle Temporal Gyrus posterior division | 23 | 0.0134 |
| cortical | Right Supramarginal Gyrus posterior division | 40 | 0.0134 |
| cerebellar | Right_VIIIb | 22 | 0.0134 |
| cortical | Left Inferior Temporal Gyrus posterior division | 29 | 0.0134 |
| cerebellar | Right_V | 4 | 0.0030 |
| cortical | Right Temporal Occipital Fusiform Cortex | 78 | 0.0030 |
| cortical | Right Inferior Frontal Gyrus pars opercularis | 12 | -0.0074 |
| cortical | Left Insular Cortex | 3 | -0.0178 |
| cortical | Left Superior Temporal Gyrus posterior division | 19 | -0.0282 |
| cortical | Right Superior Parietal Lobule | 36 | -0.0282 |
| cerebellar | Right_Dentate | 30 | -0.0282 |
| cortical | Right Inferior Temporal Gyrus temporooccipital part | 32 | -0.0491 |
| subcortical | Left Putamen | 3 | -0.0491 |
| cortical | Right Intracalcarine Cortex | 48 | -0.0491 |
| cortical | Right Occipital Fusiform Gyrus | 80 | -0.0491 |
| cortical | Left Frontal Medial Cortex | 49 | -0.0491 |
| cortical | Left Inferior Frontal Gyrus pars opercularis | 11 | -0.0595 |
| cortical | Left Parietal Operculum Cortex | 85 | -0.0595 |
| cortical | Right Occipital Pole | 96 | -0.0595 |
| cortical | Left Frontal Operculum Cortex | 81 | -0.0595 |
| cortical | Left Temporal Occipital Fusiform Cortex | 77 | -0.0699 |
| cerebellar | Right_I_IV | 2 | -0.0699 |
| subcortical | Left Thalamus | 1 | -0.0699 |
| cortical | Left Superior Temporal Gyrus anterior division | 17 | -0.0803 |
| cortical | Left Temporal Fusiform Cortex posterior division | 75 | -0.0803 |
| cerebellar | Vermis_VI | 6 | -0.0907 |
| cortical | Left Middle Temporal Gyrus anterior division | 21 | -0.0907 |
| subcortical | Right Putamen | 11 | -0.1011 |
| cortical | Right Middle Temporal Gyrus temporooccipital part | 26 | -0.1116 |
| cortical | Right Temporal Fusiform Cortex posterior division | 76 | -0.1116 |
| cortical | Right Parietal Operculum Cortex | 86 | -0.1116 |
| subcortical | Left Hippocampus | 6 | -0.1220 |
| subcortical | Right Thalamus | 9 | -0.1220 |
| cortical | Right Frontal Operculum Cortex | 82 | -0.1324 |

Table S5 – continued from previous page
| Category | Region Name | ID | Value |
| --- | --- | --- | --- |
| cortical | Right Inferior Temporal Gyrus anterior division | 28 | -0.1324 |
| subcortical | Right Caudate | 10 | -0.1428 |
| subcortical | Left Pallidum | 4 | -0.1532 |
| subcortical | Right Hippocampus | 13 | -0.1636 |
| cortical | Right Inferior Frontal Gyrus pars triangularis | 10 | -0.1741 |
| cortical | Right Parahippocampal Gyrus posterior division | 70 | -0.1741 |
| cortical | Right Superior Temporal Gyrus anterior division | 18 | -0.1741 |
| cortical | Left Subcallosal Cortex | 53 | -0.1741 |
| subcortical | Right Pallidum | 12 | -0.1741 |
| subcortical | Left Caudate | 2 | -0.1741 |
| cortical | Left Inferior Temporal Gyrus temporooccipital part | 31 | -0.1741 |
| cortical | Right Temporal Fusiform Cortex anterior division | 74 | -0.1845 |
| cerebellar | Left_I_IV | 1 | -0.1949 |
| subcortical | Right Accumbens | 15 | -0.1949 |
| cerebellar | Vermis_VIIIb | 21 | -0.1949 |
| cortical | Left Planum Temporale | 91 | -0.1949 |
| subcortical | Left Accumbens | 8 | -0.1949 |
| cortical | Left Inferior Temporal Gyrus anterior division | 27 | -0.1949 |
| cerebellar | Right_X | 28 | -0.2157 |
| cerebellar | Right_Interposed | 32 | -0.2261 |
| cerebellar | Vermis_VIIb | 15 | -0.2261 |
| cortical | Right Middle Temporal Gyrus anterior division | 22 | -0.2261 |
| cerebellar | Left_Interposed | 31 | -0.2261 |
| cerebellar | Left_X | 26 | -0.2261 |
| cerebellar | Vermis_CrusI | 9 | -0.2470 |
| cerebellar | Vermis_IX | 24 | -0.2470 |
| cortical | Right Cuneal Cortex | 64 | -0.2678 |
| cerebellar | Vermis_X | 27 | -0.2678 |
| cortical | Left Parahippocampal Gyrus posterior division | 69 | -0.2782 |
| cerebellar | Right_IX | 25 | -0.2782 |
| cortical | Right Subcallosal Cortex | 54 | -0.2886 |
| cerebellar | Left_IX | 23 | -0.2991 |
| cortical | Left Temporal Fusiform Cortex anterior division | 73 | -0.2991 |
| cortical | Left Parahippocampal Gyrus anterior division | 67 | -0.2991 |
| subcortical | Left Amygdala | 7 | -0.3199 |
| cortical | Right Planum Temporale | 92 | -0.3407 |
| subcortical | Right Amygdala | 14 | -0.3511 |
| cerebellar | Vermis_CrusII | 12 | -0.3720 |
| cortical | Right Heschl's Gyrus (includes H1 and H2) | 90 | -0.4241 |
| cortical | Right Planum Polare | 88 | -0.4449 |
| cortical | Right Parahippocampal Gyrus anterior division | 68 | -0.5074 |
| cortical | Left Planum Polare | 87 | -0.5178 |
| cortical | Left Heschl's Gyrus (includes H1 and H2) | 89 | -0.5178 |

## Notes

### Competing Interest Statement

MWM holds a US patent related to the use of CEBRA for brain decoding.

## References

1. Thomas R Insel. The nimh research domain criteria (rdoc) project: precision medicine for psychiatry. American journal of psychiatry, 171(4): 395–397, 2014.

2. René S. Kahn, Iris E. Sommer, Robin M. Murray, Andreas Meyer-Lindenberg, Daniel R. Weinberger, Tyrone D. Cannon, Michael O’Donovan, Christoph U. Correll, John M. Kane, Jim van Os, and Thomas R. Insel. Schizophrenia. Nature Reviews Disease Primers, 1(1):15067, 2015. doi: 10.1038/nrdp.2015.67.

3. Lorina Sinanaj, Konstantinos Pallis, Anahita Fazel Dehkordi, Philippe Huguelet, Stefan Kaiser, and Indrit Bègue. Mapping symptom-general and symptom-specific targets for transcranial magnetic stimulation in schizophrenia: an electric-field modeling meta-analysis. Molecular Psychiatry, pages 1–11, 2025.

4. Allyssa Chan, Andy Lu, Trisha Menon, Sabrina Wong, Kyle Valentino, Gia Han Le, Christine E. Dri, and Roger S. McIntyre. Association between negative symptoms and health-related quality of life and functional outcomes in persons with schizophrenia: A systematic review. Schizophrenia Research, 288:95–103, 2026. doi: 10.1016/j.schres.2025.12.019.

5. Silvana Galderisi, A Mucci, S Dollfus, Merete Nordentoft, P Falkai, S Kaiser, GM Giordano, A Vandevelde, Mette Ødegaard Nielsen, LB Glenthøj, et al. Epa guidance on assessment of negative symptoms in schizophrenia. European Psychiatry, 64(1):e23, 2021.

6. Saige Rutherford, Seyed Mostafa Kia, Thomas Wolfers, Charlotte Fraza, Mariam Zabihi, Richard Dinga, Pierre Berthet, Amanda Worker, Serena Verdi, Henricus G Ruhe, et al. The normative modeling framework for computational psychiatry. Nature protocols, 17(7):1711–1734, 2022.

7. Michael B First, Wayne C Drevets, Cameron Carter, Daniel P Dickstein, Lauren Kasoff, Kerri L Kim, Jonathan McConathy, Scott Rauch, Ziad S Saad, Jonathan Savitz, et al. Clinical applications of neuroimaging in psychiatric disorders. American Journal of Psychiatry, 175(9):915–916, 2018.

8. Vince D Calhoun, Stephan M Lawrie, Janaina Mourao-Miranda, and Klaas E Stephan. Prediction of individual differences from neuroimaging data. Neuroimage, 145(Pt B):135, 2017.

9. Vince D Calhoun, Robyn Miller, Godfrey Pearlson, and Tulay Adalı. The chronnectome: time-varying connectivity networks as the next frontier in fmri data discovery. Neuron, 84(2):262–274, 2014.

10. Armin Iraji, Thomas P Deramus, Noah Lewis, Maziar Yaesoubi, Julia M Stephen, Erik Erhardt, Aysneil Belger, Judith M Ford, Sarah McEwen, Daniel H Mathalon, et al. The spatial chronnectome reveals a dynamic interplay between functional segregation and integration. Human brain mapping, 40(10):3058–3077, 2019.

11. Manish Saggar, Olaf Sporns, Javier Gonzalez-Castillo, Peter A Bandettini, Gunnar Carlsson, Gary Glover, and Allan L Reiss. Towards a new approach to reveal dynamical organization of the brain using topological data analysis. Nature communications, 9(1):1399, 2018.

12. John P Cunningham and Byron M Yu. Dimensionality reduction for large-scale neural recordings. Nature neuroscience, 17(11):1500–1509, 2014.

13. Choong-Wan Woo, Luke J Chang, Martin A Lindquist, and Tor D Wager. Building better biomarkers: brain models in translational neuroimaging. Nature neuroscience, 20(3):365–377, 2017.

14. Rayan Krishnan, Pranav Rajpurkar, and Eric J Topol. Self-supervised learning in medicine and healthcare. Nature Biomedical Engineering, 6(12): 1346–1352, 2022.

15. Danilo Bzdok and Andreas Meyer-Lindenberg. Machine learning for precision psychiatry: opportunities and challenges. Biological Psychiatry: Cognitive Neuroscience and Neuroimaging, 3(3):223–230, 2018.

16. Steffen Schneider, Jin Hwa Lee, and Mackenzie Weygandt Mathis. Learnable latent embeddings for joint behavioural and neural analysis. Nature, 617(7960):360–368, 2023.

17. Steffen Schneider, Rodrigo González Laiz, Anastasiia Filippova, Markus Frey, and Mackenzie W Mathis. Time-series attribution maps with regularized contrastive learning. In Yingzhen Li, Stephan Mandt, Shipra Agrawal, and Emtiyaz Khan, editors, Proceedings of The 28th International Conference on Artificial Intelligence and Statistics, volume 258 of Proceedings of Machine Learning Research, pages 2521–2529. PMLR, 03–05 May 2025.

18. Ramprasaath R Selvaraju, Michael Cogswell, Abhishek Das, Ramakrishna Vedantam, Devi Parikh, and Dhruv Batra. Grad-cam: Visual explanations from deep networks via gradient-based localization. In Proceedings of the IEEE international conference on computer vision, pages 618–626, 2017.

19. Aditya Chattopadhay, Anirban Sarkar, Prantik Howlader, and Vineeth N Balasubramanian. Grad-cam++: Generalized gradient-based visual explanations for deep convolutional networks. In 2018 IEEE winter conference on applications of computer vision (WACV), pages 839–847. IEEE, 2018.

20. Richard SE Keefe, Terry E Goldberg, Philip D Harvey, James M Gold, Margaret P Poe, and Leigh Coughenour. The brief assessment of cognition in schizophrenia: reliability, sensitivity, and comparison with a standard neurocognitive battery. Schizophrenia research, 68(2-3):283–297, 2004.

21. Deanna M Barch, Gregory C Burgess, Michael P Harms, Steven E Petersen, Bradley L Schlaggar, Maurizio Corbetta, Matthew F Glasser, Sandra Curtiss, Sachin Dixit, Cindy Feldt, et al. Function in the human connectome: task-fmri and individual differences in behavior. Neuroimage, 80: 169–189, 2013.

22. Richard C Gershon, Molly V Wagster, Hugh C Hendrie, Nathan A Fox, Karon F Cook, and Cindy J Nowinski. Nih toolbox for assessment of neurological and behavioral function. Neurology, 80(11_supplement_3):S2–S6, 2013.

23. Alan Anticevic, Xinyu Hu, Yuan Xiao, Junmei Hu, Fei Li, Feng Bi, Michael W Cole, Aleksandar Savic, Genevieve J Yang, Grega Repovs, et al. Earlycourse unmedicated schizophrenia patients exhibit elevated prefrontal connectivity associated with longitudinal change. Journal of Neuroscience, 35(1):267–286, 2015.

24. Dae-Jin Kim, Alexandra B Moussa-Tooks, Amanda R Bolbecker, Deborah Apthorp, Sharlene D Newman, Brian F O’Donnell, and William P Hetrick. Cerebellar–cortical dysconnectivity in resting-state associated with sensorimotor tasks in schizophrenia. Human brain mapping, 41(11):3119–3132, 2020.

25. Hengyi Cao, Miklos Argyelan, Joanna Yan, Halil Aziz Velioglu, Franky Fang, Andrea Joanlanne, Simran Kang, Lara Prizgint, Jenna Schugart, Kadeem Brown, et al. Mapping cerebellar connectivity to cognition in psychosis: Convergent evidence from fmri and tms. Biological Psychiatry, 2025.

26. Sidhant Chopra, Shona M Francey, Brian O’Donoghue, Kristina Sabaroedin, Aurina Arnatkeviciute, Vanessa Cropley, Barnaby Nelson, Jessica Graham, Lara Baldwin, Steven Tahtalian, et al. Functional connectivity in antipsychotic-treated and antipsychotic-naive patients with first-episode psychosis and low risk of self-harm or aggression: a secondary analysis of a randomized clinical trial. JAMA psychiatry, 78(9):994–1004, 2021.

27. Felix Brandl, Mihai Avram, Benedikt Weise, Jing Shang, Beatriz Simões, Teresa Bertram, Daniel Hoffmann Ayala, Nora Penzel, Deniz A Gürsel, Josef Bäuml, et al. Specific substantial dysconnectivity in schizophrenia: a transdiagnostic multimodal meta-analysis of resting-state functional and structural magnetic resonance imaging studies. Biological psychiatry, 85(7):573–583, 2019.

28. Tal Geffen, Samyogita Hardikar, Jonathan Smallwood, Mariia Kaliuzhna, Fabien Carruzzo, Kerem Böge, Marco Matthäus Zierhut, Stefan Gutwinski, Teresa Katthagen, Stephan Kaiser, et al. Striatal functional hypoconnectivity in patients with schizophrenia suffering from negative symptoms, longitudinal findings. Schizophrenia Bulletin, 50(6):1337–1348, 2024.

29. Urvakhsh Meherwan Mehta, Dhruva Ithal, Neelabja Roy, Shreshth Shekhar, Ramajayam Govindaraj, Chaitra T Ramachandraiah, Nicolas R Bolo, Rose Dawn Bharath, Jagadisha Thirthalli, Ganesan Venkatasubramanian, et al. Posterior cerebellar resting-state functional hypoconnectivity: a neural marker of schizophrenia across different stages of treatment response. Biological Psychiatry, 96(5):365–375, 2024.

30. Indrit Begue, Janis Brakowski, Erich Seifritz, Alain Dagher, Philippe N Tobler, Matthias Kirschner, and Stefan Kaiser. Cerebellar and corticostriatal-midbrain contributions to reward-cognition processes and apathy within the psychosis continuum. Schizophrenia research, 246:85–94, 2022.

31. Ilaria Carta, Christopher H Chen, Amanda L Schott, Schnaude Dorizan, and Kamran Khodakhah. Cerebellar modulation of the reward circuitry and social behavior. Science, 363(6424):eaav0581, 2019.

32. Jade Awada, Farnaz Delavari, Thomas AW Bolton, Fares Alouf, Fabien Carruzzo, Noémie Kuenzi, Mariia Kaliuzhna, Tal Geffen, Teresa Katthagen, Florian Schlagenhauf, et al. A longitudinal and reproducible anti-coactivation pattern between the cerebellum and the ventral tegmental area is related to apathy in schizophrenia. Biological Psychiatry, 2025.

33. Halil Aziz Velioglu, Teresa Gomez, Juan A Gallego, Todd Lencz, Anil K Malhotra, Ariel Rokem, and Hengyi Cao. Microstructural alterations of the cerebellum-ventral tegmental area pathways in first-episode psychosis. Biological Psychiatry: Cognitive Neuroscience and Neuroimaging, 2026.

34. Narender Ramnani and Adrian M Owen. Anterior prefrontal cortex: insights into function from anatomy and neuroimaging. Nature reviews neuroscience, 5(3):184–194, 2004.

35. Human Connectome Project. HCP-EP release 1.1 reference manual. https://www.humanconnectome.org/storage/app/media/documentation/HCP-EP1.1/HCP-EP_Release_1.1_Manual.pdf. Accessed July 2026.

36. Stanley R Kay, Abraham Fiszbein, and Lewis A Opler. The positive and negative syndrome scale (panss) for schizophrenia. Schizophrenia bulletin, 13(2):261–276, 1987.

37. Brian Kirkpatrick, Gregory P Strauss, Linh Nguyen, Bernard A Fischer, David G Daniel, Angel Cienfuegos, and Stephen R Marder. The brief negative symptom scale: psychometric properties. Schizophrenia bulletin, 37(2):300–305, 2011.

38. Stefan Leucht, Myrto Samara, Stephan Heres, Maxine X Patel, Toshi Furukawa, Andrea Cipriani, John Geddes, and John M Davis. Dose equivalents for second-generation antipsychotic drugs: the classical mean dose method. Schizophrenia bulletin, 41(6):1397–1402, 2015.

39. Karen Simonyan, Andrea Vedaldi, and Andrew Zisserman. Deep inside convolutional networks: Visualising image classification models and saliency maps. arXiv preprint arXiv:1312.6034, 2013.

40. Judy Hoffman, Daniel A Roberts, and Sho Yaida. Robust learning with jacobian regularization. arXiv preprint arXiv:1908.02729, 2019.

